# α1-COP delivers sphingolipid modifiers and controls plasmodesmal callose deposition in Arabidopsis

**DOI:** 10.1101/2021.03.22.436362

**Authors:** Arya Bagus Boedi Iswanto, Minh Huy Vu, Ritesh Kumar, Jong Cheol Shon, Shuwei Wu, Da-Ran Kim, Kwak Yeon Sik, Son Geon Hui, Hobin Kang, Woe Yoen Kim, Sang Hee Kim, Kwang Hyeon Liu, Jae-Yean Kim

## Abstract

Callose is a plant cell wall polymer in the form of β-1,3-glucan, which regulates symplasmic channel size at plasmodesmata (PD). It plays a crucial role in a variety of processes in plants through the regulation of intercelluar symplasmic continuity. However, how to maintain callose homeostasis at PD in the molecular levels is poorly understood. To further elucidate the mechanism of PD callose homeostasis, we screened and identified an Arabidopsis mutant plant that exhibited excessive callose deposition at PD. Based on the Next-generation sequencing (NGS)-based mapping, other mutant allele analysis, and complementation assay, the mutated gene was shown to be *α1-COP*, which encodes a member of the COPI coatomer complex comprised of α, β, β′, γ, δ, ε, and ζ subunits. Since there is no report on the link between COPI and callose/PD, it was extremely curious to know the roles of *α1-COP* or COPI in PD regulation through callose deposition. Here, we report that loss-of-function of *α1-COP* directly elevates the callose accumulation at PD by affecting subcellular protein localization of callose degradation enzyme PdBG2. This process is linked to ERH1, an inositol phosphoryl ceramide synthase (IPCS), and glucosylceramide synthase (GCS) functions through physical interactions with the α1-COP protein. In addition, the loss-of-function of *α1-COP* also alters the subcellular localization of ERH1 and GCS proteins, results in a reduction of GlcCers and GlcHCers molecules, which are the key SL species for lipid raft formation. According to our findings, we propose that α1-COP protein, together with the SL modifiers controlling lipid raft compositions, regulates the function of GPI-anchored PD proteins and hence the callose turnover at PD and symplastic movement of biomolecules. Our findings provide the first key clue to link the COPI-mediated intracellular trafficking pathway to the callose-mediated intercellular signaling pathway through PD.

**One-sentence summary:** Plant-specific coatomer protein functions as a negative regulator of callose accumulation by regulating the translocation of sphingolipid enzymes.

## Introduction

One of the crucial components in the plant cell is callose, a polysaccharide in the form of β-1,3 glucan located at the cell walls. Callose plays a vital role in controlling the symplasmic permeability of plasmodesmata (PD) and regulates the cell-to-cell movement of signaling molecules. The callose deposition at the neck region of PD controls the symplasmic continuity. Callose is mainly synthesized by callose synthases/glucan synthase-like(s) (CalSs/GSLs) and antagonistically degraded by β-1,3-glucanases as callose degradation enzymes (BGs) (Verma and Hong, 2001; Jacobs et al., 2003; Levy et al., 2007; Barratt et al., 2011; Lee and Lu, 2011; Vaten et al., 2011; De Storme and Geelen, 2014; Iswanto and Kim, 2017; Gaudioso-Pedraza et al., 2018; Wu et al., 2018).

PD, the sophisticated symplasmic apertures, are versatile. These intracellular channels play critical roles in numerous multicellular events during plant development by conferring the molecular exchange of transcription factors, RNAs, and plant growth regulators (Zambryski and Crawford, 2000; Maule, 2008). Previous studies have described that the plasmodesmal plasma membrane (PD-PM) is distinct from common cellular PM in terms of condensed sterols and sphingolipid (SL) molecules (Grison et al., 2015; Iswanto and Kim, 2017). The enrichment of sterols and SL molecules at PM and PD-PM is often known as the membrane microdomains or lipid raft compartments (Mongrand et al., 2004; Grennan, 2007; Mongrand et al., 2010; Tapken and Murphy, 2015; Iswanto and Kim, 2017). Glycosyl inositol phosphoryl ceramide (GIPCs) and glucosyl ceramides (GlcCers) are the most abundant SL molecules found in the PM of the plant cell. Up to 64% of total sphingolipids are GIPCs, and ∼25% of the PM lipids in the *Arabidopsis thaliana* leaf are GIPCs molecules (Markham and Jaworski, 2007; Fang et al., 2016). Several studies have described the roles of lipid rafts in PD regulation, especially by regulating the subcellular localization of GPI-anchored PD proteins (Bayer et al., 2014; Grison et al., 2015; Nicolas et al., 2017; Iswanto et al., 2020).

Coat protein I (COPI) is a coatomer, a transport vesicle-bound protein complex that is responsible for various actions and several distinct secretory pathways, including ER-Golgi anterograde transport, Golgi-ER retrograde transport, intra-Golgi cargo machinery of numerous proteins and maintenance of Golgi function and structural integrity (Pepperkok et al., 1993; Gaynor et al., 1998; Schroder-Kohne et al., 1998; Paul and Frigerio, 2007; Wang et al., 2010; Ahn et al., 2015). COPI is formed of seven subunits (α/β/β’/γ/δ/ε/ζ) which are further grouped into two subcomplexes, the B-subcomplex (α/β’/ε) and F-subcomplex (β/γ/δ/ζ) (Jackson, 2014). In contrast to mammals and yeast studies, there are several isoforms of all the coatomer subunits have been identified in Arabidopsis, except γ-COP and δ-COP that contains only one isoform (Donohoe et al., 2007; Gao et al., 2014; Ahn et al., 2015; Woo et al., 2015). A previous study revealed that disruption of ε-COP subunit isoforms impairs the Golgi apparatus integrity and changes the localization of endomembrane proteins (EMPs) (Woo et al., 2015). The action of COPI within intracellular trafficking is tightly connected to the cargo molecules containing the dilysine KKXX and KXKXX motifs presented on their C-terminal tail (Schroder-Kohne et al., 1998; Eugster et al., 2004; Jackson et al., 2012; Ma and Goldberg, 2013). Also, the recent study of Arabidopsis *α-COP* reveals that *α2-COP* is essential for plant growth and development by maintaining the morphology of the Golgi apparatus through the subcellular localization of a protein harboring dilysine motif, p24δ5 (Gimeno-Ferrer et al., 2017).

Previously, we generated dexamethasone inducible *RNAi* line of *Glucan synthase-like 8* (*dsGSL8 RNAi*), which is defective in tropism due to the absence of GSL8-induced callose deposition (Han et al., 2014). In this study, EMS mutagenesis of *dsGSL8 RNAi* resulted in a mutant that rescued the tropic responses and showed a high PD callose phenotype in Arabidopsis hypocotyls. In this mutant line, the NGS-based mapping (NGM) revealed a point mutation in the AT1G62020 (*α1-COP*) with single amino acid substitution. We also found that other T-DNA inserted *α1-COP* mutants exhibited excessive callose phenotype. Moreover, in the *α1-cop* mutant, the subcellular localization of ERH1 and GCS, which are two SL pathway enzymes with dilysine motif, were mislocalized. We also found that the localization of PdBG2, one of the callose degrading GPI-anchored enzymes, was also altered in the *α1-cop* mutant. Here, we provide evidence for the novel function of α1-COP in regulating PD callose deposition through SL modifiers cargo machinery.

## Results

### Loss-of-function of *α1-COP* exhibits excessive callose accumulation

We treated tropism defective *dsGSL8 RNAi* plants (Han et al., 2014) with ethyl methanesulfonate (EMS), strikingly we found several *dsGSL8 RNAi*-EMS lines displayed rescued tropism responses (data not shown). Furthermore, we selected one mutant line for subsequent analyses. NGM result predicted the point mutation in the AT1G62020 (*α1-COP*), with a single amino acid substitution, *α1-cop-4* (G486D substitution) **(Supplemental Figure 1A, Figure 1A, B)**. For further study, we characterized two more mutants of *α1-COP*, *α1-cop-1* (SALK_078465) and *α1-cop-5* (SALK_003425), which have the T-DNA insertion at different positions of the gene and shows different transcript abundance **(Figure 1C, D)**. Phototropic response by Arabidopsis hypocotyl is associated with callose dependent modulation of PD permeability (Han et al., 2014). Hence, we conducted a PD permeability assessment using the HPTS movement assay for these mutants. The HPTS movement analysis was conducted at 3-day-etiolated seedlings of wild-type Col-0 and *α1-cop* mutant plants. Interestingly, three independent alleles of *α1-cop* mutants show reduced HPTS diffusions in comparison to wild-type Col-0 **(Supplemental Figure 1B, C).** To test whether this PD permeability alteration is linked to the callose accumulation or not, aniline blue staining was done to check the callose level in etiolated seedlings and rosette leaves of Arabidopsis wild-type Col-0 and *α1-cop* mutant plants. Interestingly, *α1-cop* mutant plants shown elevated callose levels in both hypocotyl and Arabidopsis rosette leaves **(Figure 1E, F, G, H)**.

**Figure 1.**
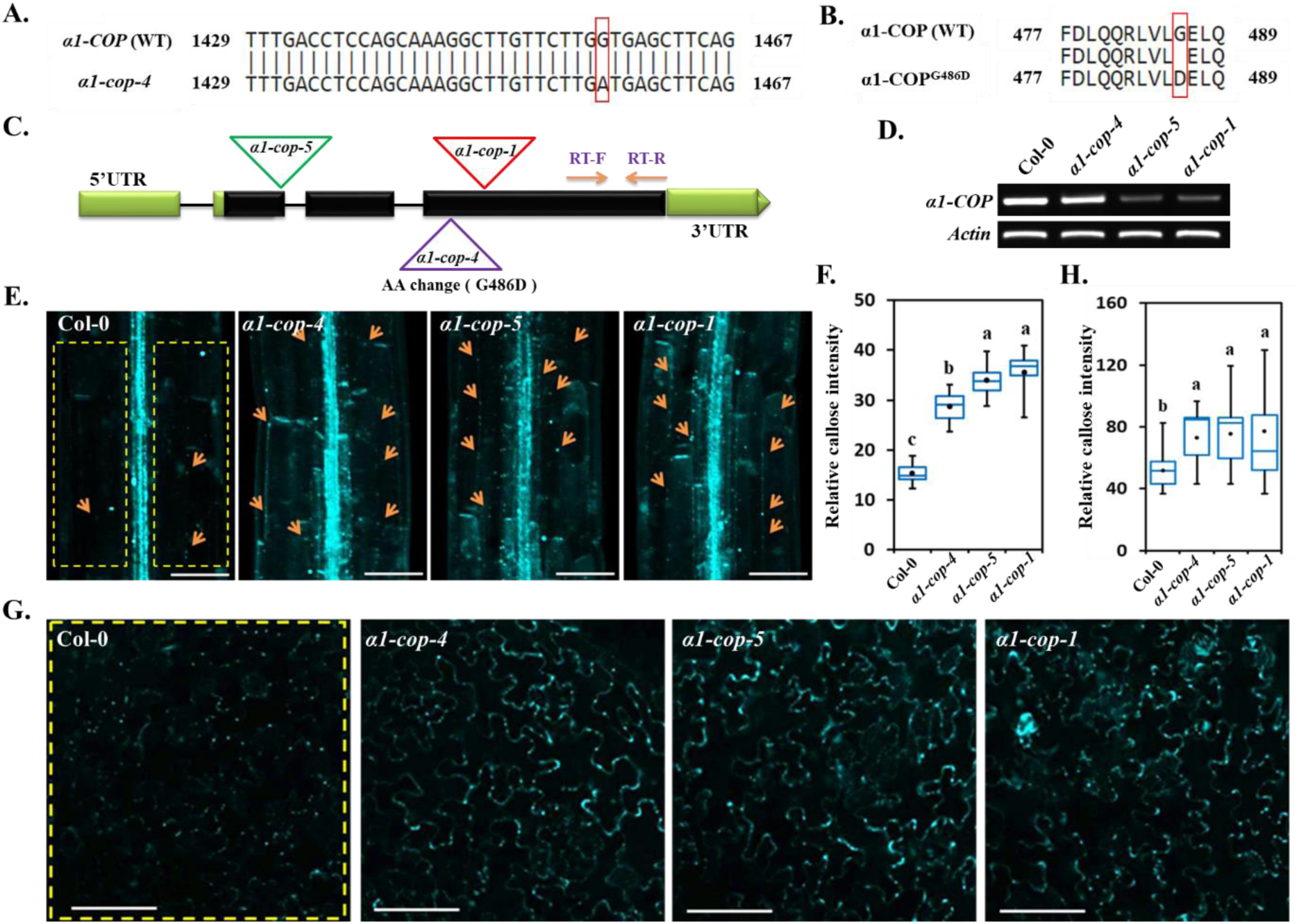
Callose accumulation is increased in the *α1-cop* mutants. **(A)** Nucleotide sequence alignment of *α1-COP* WT (wild-type Col-0) and *α1-COP* EMS mutant. The red box indicates single nucleotide mutation (G to A). **(B)** Amino acid (AA) sequence alignment of α1-COP WT and α1-COP^G486D^ (EMS mutant). The red box indicates single AA mutation (G to D). **(C)** Gene structure of *α1-COP* with three allele mutations. *α1-cop-4* was generated from EMS mutation and identified by NGS-based mapping. *α1-cop-1* and *α1-cop-5* were collected from ABRC as T-DNA insertions. **(D)** RT-PCR analysis from wild type and *α1-cop* mutants. One pair primer was designed at the C-terminal domain close to 3’UTR region as shown in the figure **(A)**. **(E)** Callose deposition analysis of Arabidopsis hypocotyls from wild-type Col-0 and *α1-cop* mutants. Scale bars: 100 µm. **(F)** Relative callose intensity quantification of Arabidopsis hypocotyls (*n=*10). **(G)** Callose deposition analysis of Arabidopsis rosette leaves. Scale bars: 100 µm. **(H)** Relative callose intensity quantification of Arabidopsis rosette leaves (*n=*15). All 3 independent biological experiments were performed and statistical significances were done by One-Way ANOVA with Tuckey-Kramer test. Yellow square dotted lines **(E, G)** were designated as region of interest (ROI) for measuring signal intensity.

We also generated transgenic plants overexpressing *α1-COP* (*p35S::α1-COP)* in the wild-type Col-0 background. α1-COP-OE#2 and α1-COP-OE#3 were selected **(Supplemental Figure 2A)** and subsequently tested for PD permeability analysis. *α1-COP* overexpression plants exhibited more substantial HPTS diffusion and less callose accumulation as compared with wild-type Col-0 **(Supplemental Figure 2B-E)**. We also checked the phototropism phenotype from *α1-cop* mutants and *α1-COP* overexpression plants. Consistent with our initial mutant screening, three independent mutant alleles of *α1-COP* showed faster phototropism and increased curvature angle. Conversely, the hypocotyl curvature angle was attenuated in the *α1-COP* overexpression plant in comparison to the wild-type Col-0 and *GSL8* overexpression plants **(Supplemental Figure 1D, E)**. In Arabidopsis, there are two isoforms of α-COP proteins, and it had been reported that α2-COP protein is critical for plant growth and development by regulating the secretory pathway of p24δ5 protein as well as maintaining the morphology of Golgi apparatus (Gimeno-Ferrer et al., 2017). Next, to anticipate any role of *α2-COP* in controlling the callose-mediated PD permeability, the callose staining assay was done to determine the callose level in hypocotyls of *α2-cop* mutants. Surprisingly, two independent *α2-cop* mutant alleles and wild-type Col-0 hypocotyls showed a similar level of callose deposition **(Supplemental Figure 3A-C)**, and the phototropism was comparable to wild-type Col-0 plants **(Supplemental Figure 3D, E)**, indicating that *α2-COP* is not involved in the callose-induced phototropism. In short, our results suggest that *α1-COP* functions specifically to increase PD permeability by decreasing callose accumulation in Arabidopsis.

### α1-COP is a *trans*-Golgi-localized protein, but partially localized at PD

α1-COP is a member of COPI that facilitate retrieval/retrograde transport of numerous proteins from Golgi to endoplasmic reticulum (ER), and for intra-Golgi delivery (Paul and Frigerio, 2007; Ahn et al., 2015). Therefore, to determine if in plants also α1-COP localized at Golgi compartment, α1-COP fusion proteins with GFP/RFP-tag at either N-terminal or C-terminal positions were generated and transiently expressed in *Nicotiana benthamiana* epidermal cells **(Figure 2A).** All the configurations and tags showed a similar fluorescent pattern **(Figure 2A)**. Next, we expressed well-known *trans-*Golgi protein, ERH1 (Wang et al., 2008), along with α1-COP, and found α1-COP was highly co-localized with ERH1 **(Figure 2B)**. Moreover, these signals also showed PD-like punctate fluorescent signals at the cell periphery. To further verify the PD localization, we examined the co-localization of α1-COP and aniline blue-stained PD callose. Aniline blue is a widely used PD marker that stains callose localized at the PD neck (Vaten et al., 2011). The transiently expressed α1-COP signals partially co-localized with aniline blue at PD in *N. benthamiana* **(Figure 2C)**. The Arabidopsis transgenic plant overexpressing α1-COP (p35S::GFP:α1-COP) were generated in the wild-type Col-0 background to determine the subcellular localization of α1-COP. Consistent with transient expression data, stable GFP:α1-COP and aniline blue signals were partially co-localized at PD **(Figure 2D)**. Together with the colocalization data with the other PD marker, PDLP5, PDLP1 and PdBG2 (shown in **Figure 3C, Figures 10A, C**), these results suggest that α1-COP is a *trans-*Golgi-localized protein that also partially located at PD.

**Figure 2.**
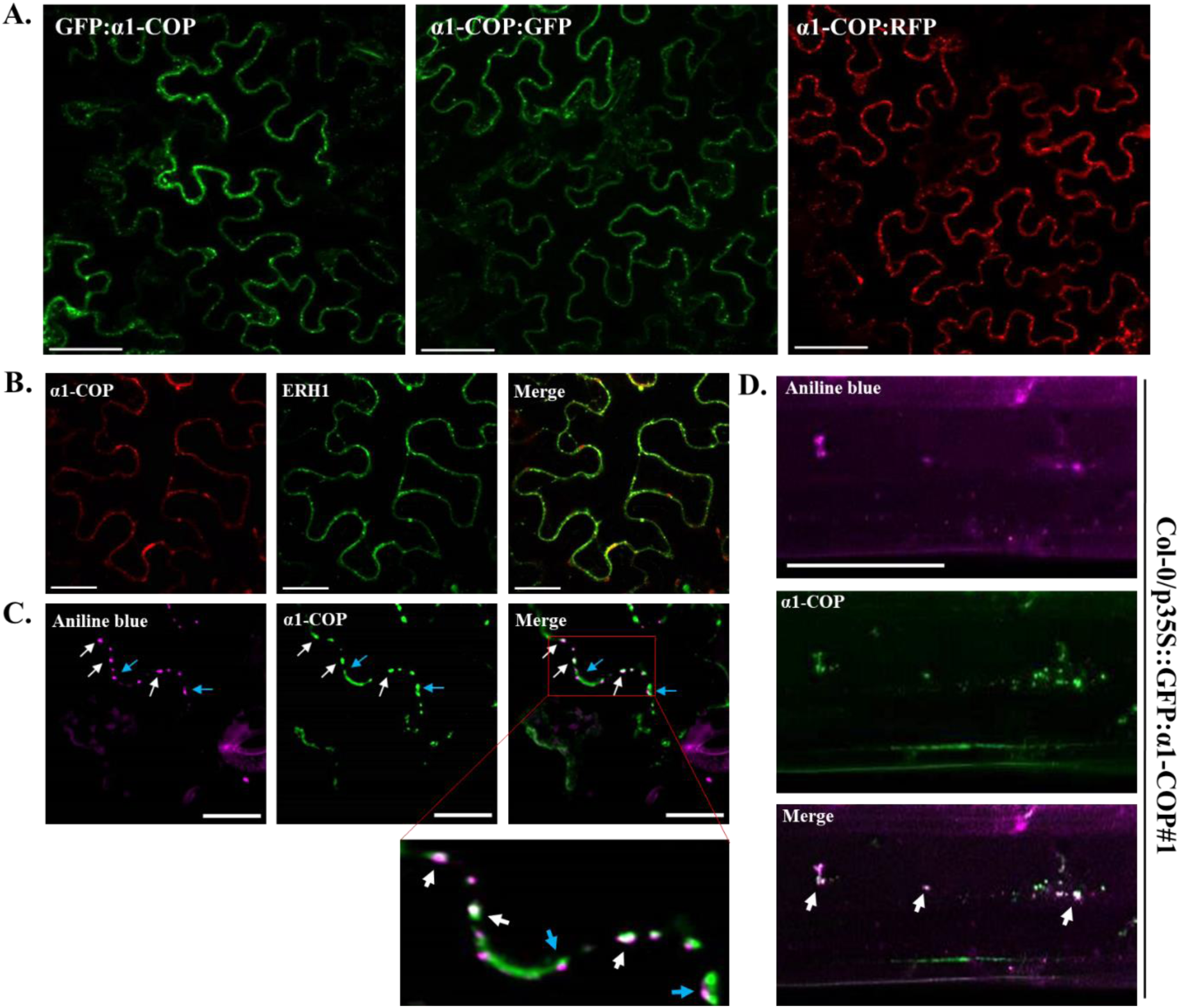
α1-COP is localized at *trans*-Golgi compartment and partially localized at PD. **(A)** Confocal images of *N. benthamiana* epidermal cells transiently expressing 3 different fusion proteins of α1-COP. Both GFP and RFP signals were detected at the cell periphery and exhibited PD-like punctate localization. Scale bars: 50 µm. **(B)** Confocal images of *N. benthamiana* epidermal cells transiently expressing α1-COP:RFP and ERH1:GFP (*trans*-Golgi protein). Merge picture with green and red signals showed that α1-COP:GFP is localized to *trans*-Golgi compartment. Scale bars: 50 µm. **(C)** Transient expression of GFP:α1-COP and chemical staining of callose using aniline blue. The white arrows illustrate co-localization at PD, whereas the blue arrows illustrate the GFP signals do not co-localize with Aniline blue signal. Scale bars: 20 µm. **(D)** Arabidopsis hypocotyl transgenic plant expressing GFP:α1-COP stained with aniline blue. The white arrows illustrate co-localization at PD. Scale bars: 50 µm.

**Figure 3.**
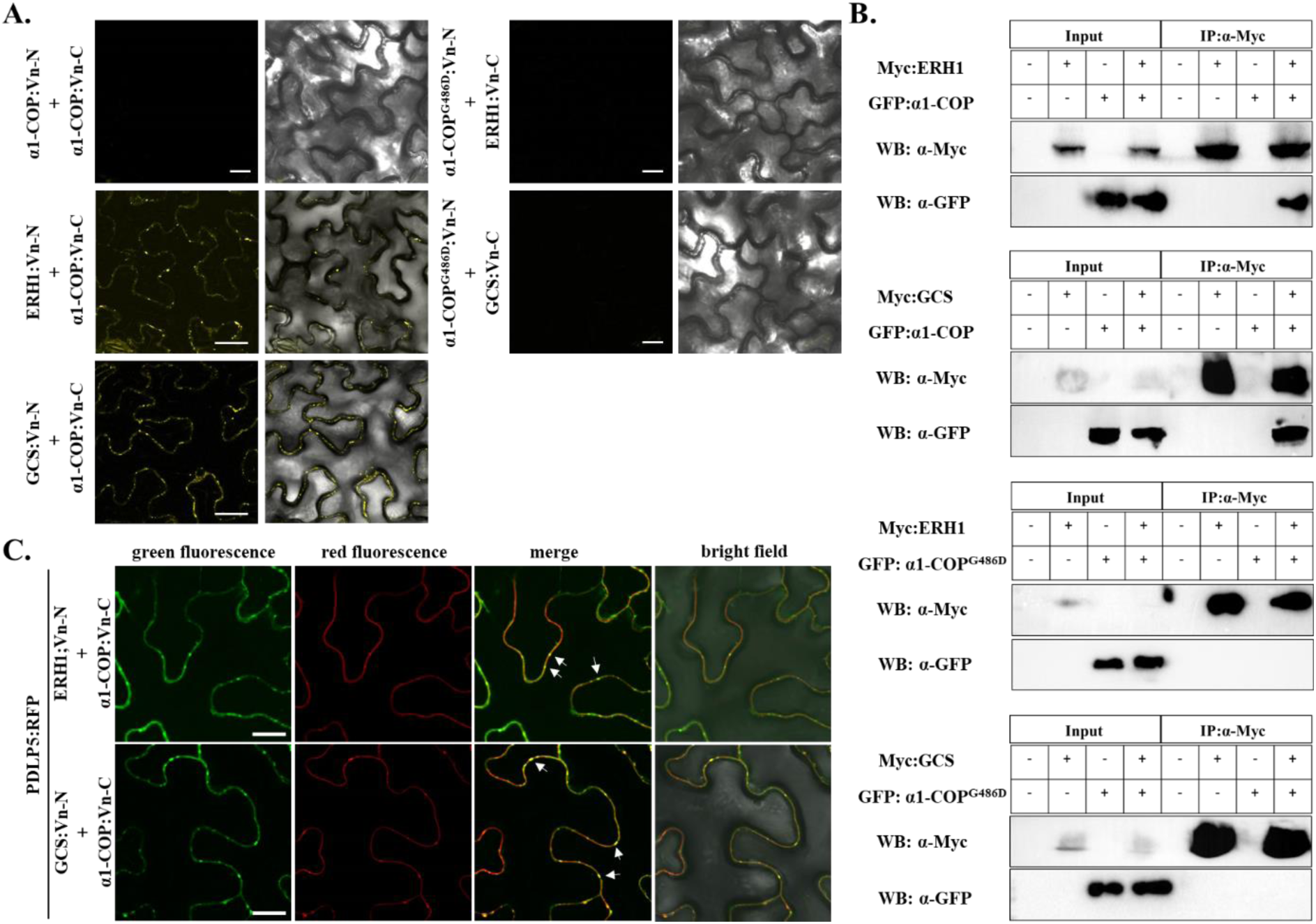
α1-COP interacts with ERH1 and GCS. **(A)** BiFC assay of the interactions between α1-COP and ERH1, α1-COP and GCS, α1-COP^G486^ and ERH1, α1-COP^G486^ and GCS. Various combinations of BiFC vectors were transiently expressed in *N. benthamiana* leaves. The infiltrated leaves were subjected for confocal imaging 3-days post infiltration. Three biological replicates were performed for each sample. Scale bars = 50 µm. **(B)** Co-IP analysis of the interaction between α1-COP and ERH1, α1-COP and GCS, α1-COP^G486^ and ERH1, α1-COP^G486^ and GCS. Various combinations of Myc-ERH1, Myc-GCS, GFP-α1-COP and GFP-α1-COP^G486^ fusion proteins as indicated were transiently expressed in *N. benthamiana* leaves followed by IP using protein A agarose bead. A GFP or Myc antibody was used to detect the proteins. **(C)** α1-COP, ERH1 and GCS interactions are partially located at PD. Confocal images of *N. benthamiana* leaves agro-infiltrated with BiFC constructs of ERH1:Venus-N, GCS:Venus-N, and α1-COP:Venus-C (co-expressed with PDLP5-RFP). Green fluorescence resulting from the interactions of ERH1:Venus-N and α1-COP:Venus-C, GCS:Venus-N and α1-COP:Venus-C were observed at the cell periphery. The green spots co-localize with the red fluorescent signals emitted by RFP-labeled PDLP5 (white arrows). Three biological replicates were performed for each sample. Scale bars: 20 µm.

### α1-COP interacts with sphingolipid modifiers ERH1 and GCS

To examine the role of the α1-COP protein in the formation of a heptameric protein complex coatomer on vesicles, we first used bimolecular complementation (BiFC) to confirm the interactions of the coatomer complex components *in planta*. β’2-COP and ε1-COP were selected from the B-subcomplex members, and δ-COP was preferred from the F-subcomplex members. BiFC analysis clearly showed the α1-COP interaction with ε1-COP, β’2-COP, and δ-COP **(Supplemental Figure 4)**. The primary function of COPI vesicles is to transport proteins and lipids back to the previous compartment along the secretory pathway. Furthermore, transmembrane proteins containing a KKXX or a KXKXX motif on their C-terminal tail are COPI dependent cargo machinery, which is retrieved from the Golgi apparatus to the ER (Spang, 2013). Several studies in mammals shown that the COPI specifically interacts with SL species (Chaudhary et al., 1998; Contreras et al., 2012). In Arabidopsis, several well-known SL enzymes such as LONGEVITY ASSURANCE GENE ONE HOMOLOGs (LOHs), ERH1, and GCS (Wang et al., 2008; Msanne et al., 2015; Xie et al., 2015; Iswanto et al., 2020) have been reported.

To determine if α1-COP also interacts with SL enzymes *in planta*, Arabidopsis SL enzymes were analyzed for the presence of dilysine motif at their C-terminal domain. Detail analysis showed that most of the SL pathway enzymes in Arabidopsis have dilysine motifs, and subsequently, we selected two prominent SL enzymes ERH1 and GCS for further study **(Supplemental Figure 5A)**. To examine the function of α1-COP at ERH1 and GCS cargo molecules, BiFC assay was performed. The BiFC assay showed that α1-COP interacts with ERH1 and GCS **(Figure 3A)**. We further confirmed these interactions using co-immunoprecipitation (Co-IP) analyses. GFP:α1-COP with or without Myc:ERH1/Myc:GCS was transiently expressed in *N. benthamiana* leaves. Co-IP followed by immunoblot analyses exhibited that α1-COP interacts with ERH1 and GCS **(Figure 3B)**. The fluorescent signals from the both BiFC assay using α1-COP and ERH1/GCS were detected at the entire cell periphery along with some PD-like punctate spots. Further, we validated the PD localization by transiently expressing two sets of BiFC constructs in *N. benthamiana;* (ERH1:nVenus and α1-COP:cVenus, GCS:nVenus and α1-COP:cVenus) along with PLASMODESMATA-LOCATED PROTEIN 5 (PDLP5:RFP) (Thomas et al., 2008). Interestingly, GFP punctate spots on the cell periphery showed perfect co-localization with PDLP5:RFP indicating PD localization of α1-COP interactions **(Figure 3C)**. These results confirm that α1-COP interacts with SL enzymes, ERH1, and GCS at the cell periphery along with PD.

### Single amino acid substitution of α1-COP affects its interaction with ERH1 and GCS

In yeast, a single amino substitution at N-terminal domain of α1-COP have shown several defective phenotypes in the intracellular transports of dilysine cargo molecules (Schroder-Kohne et al., 1998; Eugster et al., 2000; Kim et al., 2011). Next, to determine the effect of the α1-COP^G486D^ single amino substitution mutant at ERH1 and GCS interactions, EMS mutated α1-COP^G486D^ was amplified and cloned. Firstly, the mutant version of α1-COP was fused to GFP (p35S::GFP:α1-COP^G486D^) **(Supplemental Figure 5B)** to check the subcellular localization. The GFP:α1-COP^G486D^ and α1-COP:RFP were transiently co-expressed in *N. benthamiana*. Confocal images showed that α1-COP^G486D^ was highly co-localized with wild type α1-COP **(Supplemental Figure 5C)**. This result indicates that a single amino substitution (G486D) at N-terminal of α1-COP does not change its subcellular localization.

Next, BiFC assay was performed to analyze the effect of single amino substitution on the α1-COP^G486D^ interactions with ERH1 and GCS. Two sets of constructs (ERH1:nVenus and α1-COP^G486D^cVenus, GCS:nVenus and α1-COP^G486D^:cVenus) were transiently expressed in *N. benthamiana* leaves separately. Surprisingly, no interactions with ERH1 or GCS were observed by BiFC **(Figure 3A)**. We further validated the BiFC data using Co-IP assay. GFP:α1-COP^G486D^ with or without Myc:ERH1/Myc:GCS were transiently expressed in *N. benthamiana* leaves. Co-IP followed by immunoblot analyses showed that α1-COP does not interact with ERH1 and GCS **(Figure 3B)**. The BiFC data agree with Co-IP data, validating that α1-COP^G486D^ does not interact with ERH1 and GCS. Together, these data suggest that glycine residue at 486 aa in α1-COP plays a critical role in maintaining the physical interactions with ERH1 and GCS SL pathway enzymes. However, this mutation does not affect the subcellular localization of α1-COP.

### Loss of function of *α1-COP* alters the subcellular localization of ERH1 and GCS

Previously it was reported that the reduction in the COPI complex had favored the mislocalization of cholesterol, sphingolipids, Rac1, and Cdc42 away from the plasma membrane into the cytoplasmic compartment in the animal system (Misselwitz et al., 2011). Similarly, the cellular localization of AtERH1 in the absence of α1-COP was tested. As α1-COP, partially localized at PD, there may be a probability that its interacting SL modifier ERH1 also confines to similar cellular loci. To check this possibility ERH1:GFP and PDLP5:RFP were transiently co-expressed in *N. benthamiana.* Similar to α1-COP, ERH1:GFP fluorescent signal was partially co-localized with PDL5:RFP signals at PD **(Figure 4A)**. Also, the stable transgenic plants overexpressing ERH1:GFP in the wild-type Col-0 and *α1-cop-5* plants were generated. Consistent with transient expression, wild-type Col-0 plants overexpressing ERH1:GFP showed PD-like peripheral punctate spots **(Figure 4B-D, Supplemental Figure 6A)**, but surprisingly ERH1 was mislocalized in the absence of α1-COP protein, it lacked the PD-like punctate spots and was mainly accumulated in the cytoplasm **(Figure 4E, Supplemental Figure 6B)**. Also, peripheral punctate spots shown by ERH1:GFP in the wild-type Col-0 plants were strongly co-localized with aniline blue signals at PD **(Figure 4F)**, which further validated the PD localization of ERH1. In contrast, cytoplasmic ERH1:GFP signals in the *α1-cop-5* plant did not co-localize with aniline blue at PD **(Figure 4G)**. Furthermore, to investigate the roles of α1-COP and ERH1 in intracellular trafficking, we used Brefeldin A (BFA) which is known to block protein transport between the endoplasmic reticulum (ER) and the Golgi apparatus (Geng et al., 2015). Previously, in the absence of BFA, GFP signals of α1-COP **(Figure 2C)** and ERH1 **(Figure 4A**) showed PD punctate spots. However, *N. benthamiana* leaves transiently expressing GFP:α1-COP or ERH1:GFP infiltrated with BFA (6 h before observation) resulted in the accumulation of massive fluorescent aggregates at cytoplasm **(Supplemental Figure 7)**. These results indicate that α1-COP and ERH1 are involved in intracellular trafficking, and the absence of α1-COP interferes subcellular localization of ERH1.

**Figure 4.**
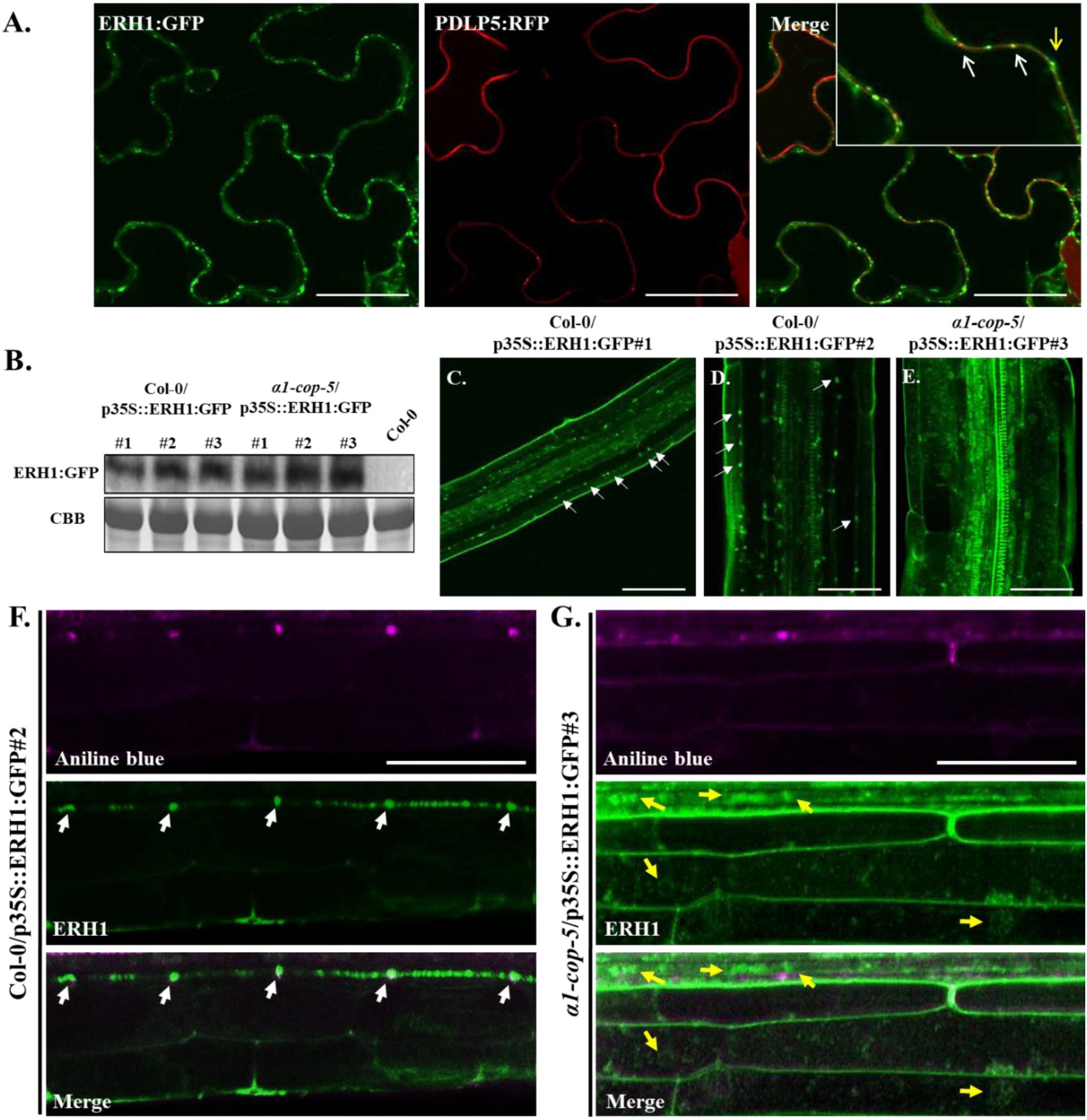
Plasmodesmata localization of ERH1 is altered in the *α1-cop-5* mutant. **(A)** ERH1 and PDLP5 is partially co-localized at PD. Confocal images of *N. benthamiana* epidermal cells transiently expressing ERH1:GFP and PDLP5:RFP. White arrows indicate co-localization signals from GFP and RFP, whereas yellow arrow indicates GFP signals do not co-localize with RFP signals. Scale bars: 50 µm. **(B)** Western blot analysis of ERH1 protein expressed in the wild-type Col-0 and *α1-cop-5* mutant plants. **(C)** Arabidopsis primary root of wild-type Col-0 plant expressing ERH1:GFP. The GFP signals in specific punctate spots are indicated by white arrows. Scale bars: 80 µm. **(D)** Arabidopsis primary root of wild-type Col-0 plant expressing ERH1:GFP. The GFP signals in specific punctate spots are indicated by white arrows. Scale bars: 30 µm. **(E)** Arabidopsis primary root of *α1-cop-5* mutant expressing ERH1:GFP. Scale bars: 30 µm. **(F)** GFP-tagged ERH1 signals are partially co-localized with aniline blue-stained callose in the primary root of Arabidopsis transgenic plant (wild-type Col-0 background). White arrows illustrate GFP signals co-localized with aniline blue signals (magenta) at PD. Scale bar: 40 µm. **(G)** GFP-tagged ERH1 signals do not co-localize (yellow arrows) with aniline blue-stained callose (magenta) in the primary root of Arabidopsis transgenic plant (*α1-cop-5* mutant background). Scale bar: 40 µm.

GCS plays a vital role in plant growth and development. Previous studies showed that GCS is localized at the ER compartment (Melser et al., 2010; Msanne et al., 2015). Consistent with previous findings, GCS:GFP was highly co-localized with ER retention signal HDEL:RFP **(Figure 5A),** along with some PD-like punctate spots at the cell periphery **(Figure 5A)**. To check whether GCS and ERH1 share the typical subcellular localization, ERH1:GFP and GCS:RFP were transiently co-expressed in *N. benthamiana* and noticeably GCS:RFP was partially co-localized with ERH1:GFP **(Figure 5B).** To verify PD-like peripheral punctate spots showed by GSC:GFP are located at PD, transiently expressed GCS:GFP tobacco leaves were stained with aniline blue. GCS:GFP exhibited a good co-localization with aniline blue-stained callose at PD **(Figure 5C-D)**. To validate that GCS is partially localized at PD, transgenic plants overexpressing GCS:GFP were generated. Consistent with transient expression, GCS:GFP in the wild-type Col-0 background showed partial co-localization with aniline blue signals at PD **(Figure 5E)**. However, in the absence of α1-COP protein, GCS was mainly localized at cytoplasm and did not co-localize with PD callose **(Figure 5F)**. Taken together, these results suggest that α1-COP protein directly or indirectly modulates the subcellular localization of ERH1 and GCS SL modifiers in Arabidopsis.

**Figure 5.**
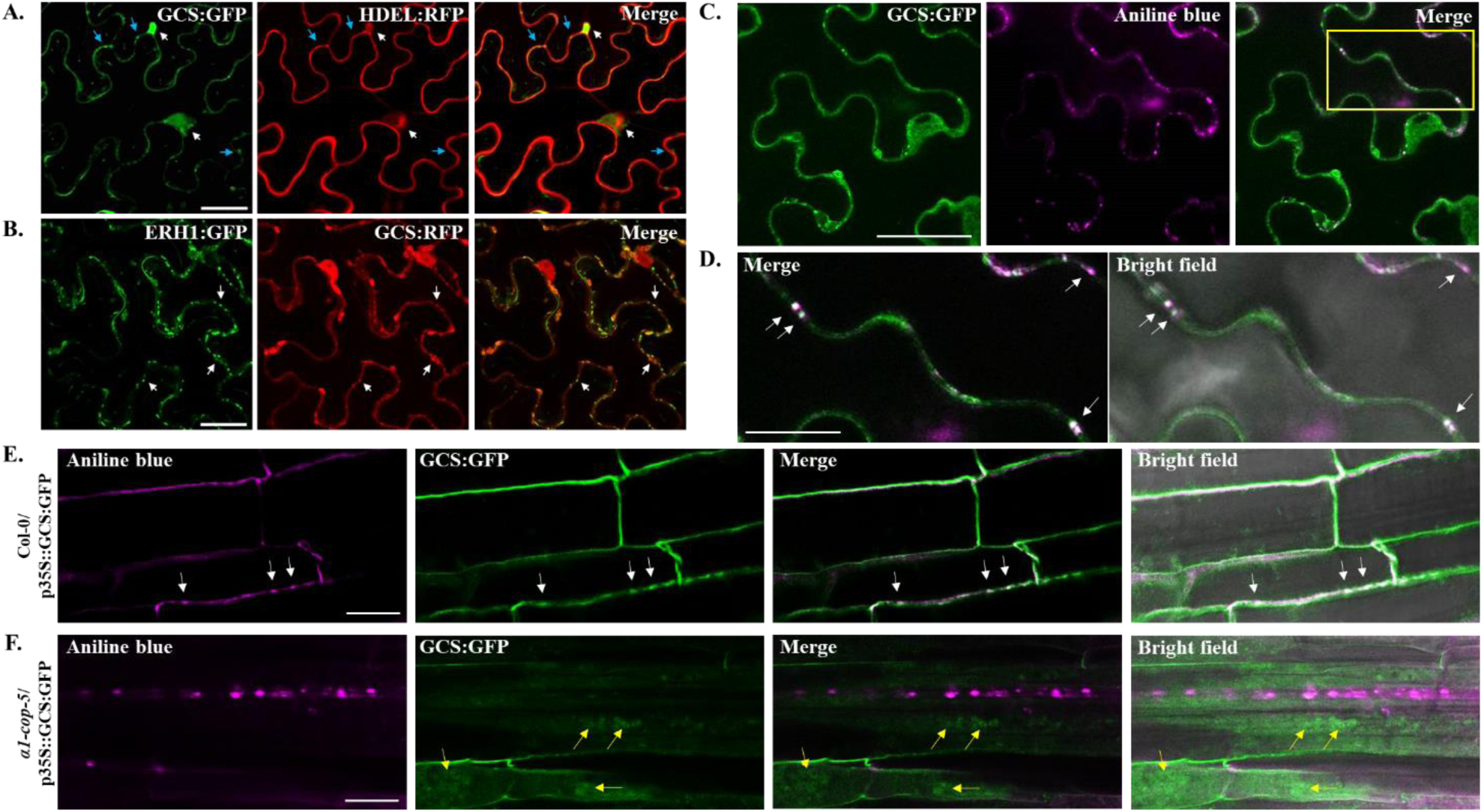
Subcellular localization of GCS is altered in the *α1-cop-5* mutant. **(A)** *N. benthamiana* epidermal cells transiently expressing fluorescent fusion proteins of GCS:GFP and HDEL:RFP. White arrows indicate co-localization of GFP and RFP signals at ER. Blue arrows indicate GFP signals do not co-localize with RFP signals. Scale bars: 50 µm. **(B)** Confocal images of *N. benthamiana* epidermal cells transiently expressing ERH1:GFP and GCS:RFP. White arrows indicate co-localization of GFP and RFP signals. Scale bars: 50 µm. **(C)** Confocal images of *N. benthamiana* epidermal cells transiently expressing GCS:GFP. Sample was stained with aniline blue for PD localization analysis. Scale bar: 50 µm. **(D)** Confocal images of *N. benthamiana* epidermal cells transiently expressing GCS:GFP **(C)** in yellow square line. White arrows indicate co-localization of GFP and magenta (aniline blue) signals at PD. Scale bar: 20 µm. **(E, F)** Transgenic GCS:GFP Arabidopsis wild-type Col-0 **(E)** and *α1-cop-5* mutant **(F)** showing fluorescence as punctate spots on the cell walls and in cytoplasm of primary root cells, respectively. White arrows indicate the GFP signals co-localize with aniline blue signals (magenta) at PD. Yellow arrows indicate the GFP signals do not co-localize with aniline blue signals (magenta) at PD Scale bar: 20 µm.

### *α1-COP* is involved in the sphingolipid biosynthesis pathway

Since ERH1 and GCS interacted with α1-COP, and loss of function of *α1-COP* altered their subcellular localizations, we hypothesized that SL compositions are associated with α1-COP function. To experimentally test the hypothesis, firstly, the transcript level of *ERH1* and *GCS,* along with several genes involved in the SL pathway was analyzed. In the absence of *α1-COP*, transcript levels of *ERH1* and *GCS* were similar to that of wild-type Col-0 plants. In contrast, plants overexpressing *α1-COP,* strongly induced the expression level of several SL pathway genes as compared to wild-type Col-0 plants **(Supplemental Figure 8)**. Next, the SL molecules reported in plants such as LCBs, ceramides, hydroxyceramides, GlcCers, GlcHCers and GIPCs (Markham et al., 2006; Magnin-Robert et al., 2015; Ali et al., 2018; Yan et al., 2019; Iswanto et al., 2020) were analyzed from wild-type Col-0, *α1-cop-1*, *α1-cop-5*, α1-COP-OE#1 and α1-COP-OE#2 overexpression plants. The SL profiling exhibited significant alterations in the ceramides, GlcCers, and GlcHCers levels as compared to wild-type Col-0 **(Figure 6)**. Loss of function of *α1-COP* showed a significant reduction in the total ceramides, GlcCers, and GlcHCers contents, conversely α1-COP overexpression plants displayed significant elevation in the several molecules of GlcCers **(Figure 6)**. Overall, these results indicate that *α1-COP* is mainly involved in the GlcCers and GlcHCers homeostasis maintenance presumably through intracellular regulation of SLs modifiers in Arabidopsis.

**Figure 6.**
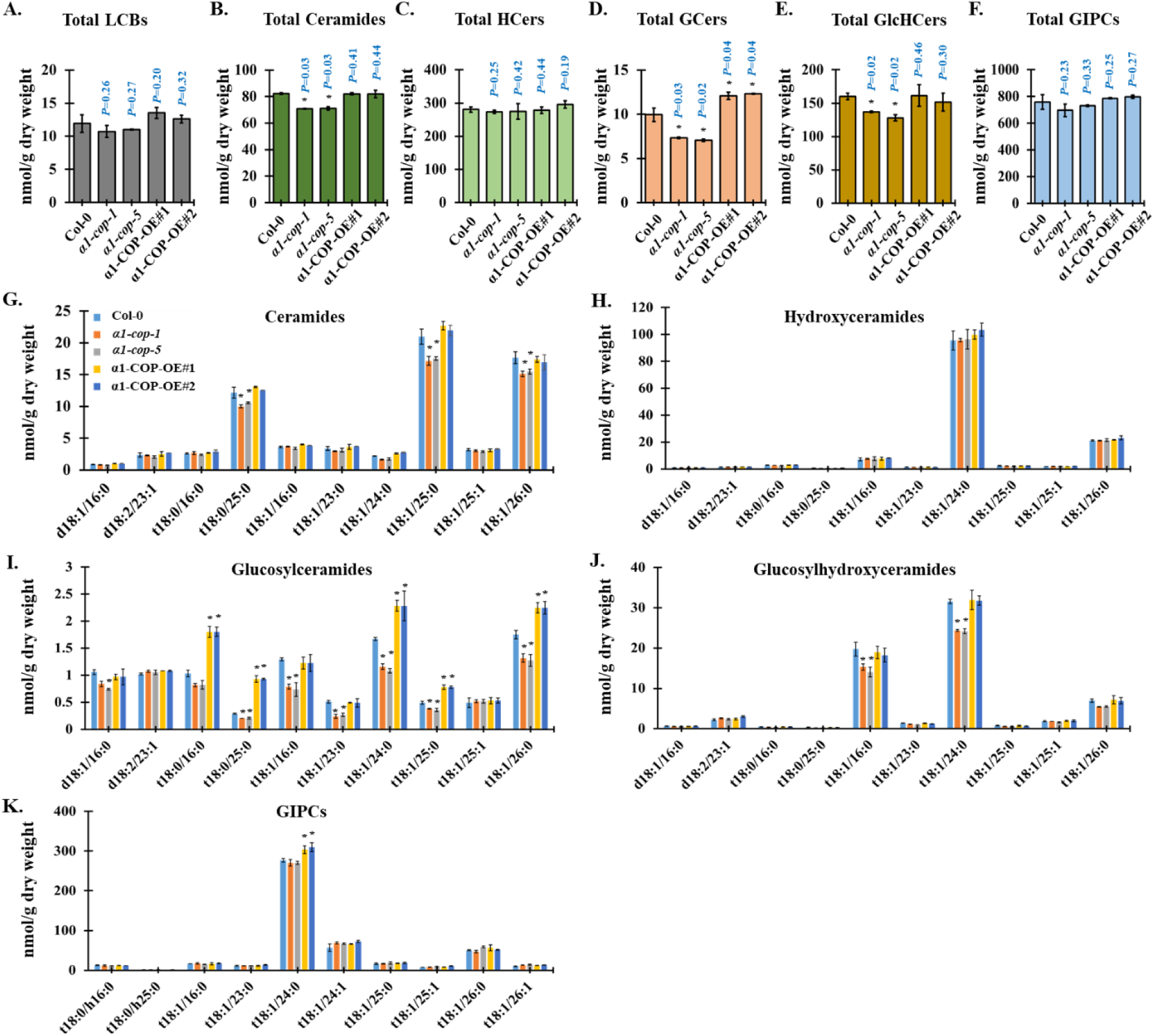
Loss-of-function of *α1-COP* reduces ceramides, GlcCers and GlcHCers. **(A-F)** Measurement of SLs from wild-type Col-0, *α1-cop-1*, *α1-cop-5*, α1-COP-OE#2 and α1-COP-OE#3 plants included total LCBs **(A)**, total ceramides **(B)**, total hydroxyceramides **(C)**, total GlcCers **(D)**, total GlcHCers **(E)** and total GIPCs **(F)**. (G-K) SLs species characterized by LCB (d18:0, d18:1, d18:2, t18:0 and t18:1) and fatty acid (FA) (16:0–26:1) from wild-type Col-0, *α1-cop-1*, *α1-cop-5*, α1-COP-OE#1 and α1-COP-OE#2 overexpression plants included ceramides **(G)**, hydroxyceramides **(H)**, GlcCers **(I)**, GlcHCers **(J)** and GIPCs **(K)**. Measurements are the average of four independent biological experiments (*n* = 200). Data are means ± s.d. Statistical significances were done by two-tailed Student’s t-test; *\*P* < 0.05, *\*\*P* < 0.01.

### Loss of function of *α1-COP* changes the subcellular localization of PdBG2

As the loss of function of the *α1-COP* mutant showed enhanced PD callose phenotype, a question arises whether the subcellular localization of GPI-anchored proteins was perturbed in the *α1-cop* mutants. Previous studies have remarkably identified the effect of sterols along with SL alteration in the functions of GPI-anchored PD proteins, PdBG2 and PDCB1 (Farquharson, 2015; Grison et al., 2015; Iswanto and Kim, 2017; Iswanto et al., 2020). PdBGs belong to the group of GPI-anchored PD proteins, which are highly linked to sterol and SL-enriched lipid raft in *planta*. In Arabidopsis, two PdBGs (PdBG1 and PdBG2) have been well studied, which are known to degrade callose deposition at the neck region of PD (Zavaliev et al., 2011; Benitez-Alfonso et al., 2013; Zavaliev et al., 2013; Zavaliev et al., 2016; Yeats et al., 2018). Firstly, transgenic plants overexpressing GFP:PdBG2 in the wild-type Col-0 and *α1-cop*-*5* plants were generated. Furthermore, the subcellular localization of PdBG2 in the Arabidopsis cotyledons and hypocotyls was analyzed in the mutant and wild-type Col-0 backgrounds. Consistent with previous reports (Iswanto et al., 2020), GFP:PdBG2 was localized at PD in the wild-type Col-0 background **(Figure 7A, C).** In contrast, in *α1-cop-5*, GFP:PdBG2 failed to accumulate at PD **(Figure 7B, D).** To confirm the mislocalization of PdBG2 in the *α1-cop-5* mutant, the cotyledons were stained with aniline blue before imaging. As expected, the observed GFP fluorescence in wild-type Col-0 plants showed perfect co-localization with PD callose, whereas GFP fluorescence depicted in the *α1-cop-5* did not co-localize with PD callose **(Figure 7A, B)**. The PdBG2 localization was also analyzed in Arabidopsis hypocotyls. A similar mislocalization event was observed in hypocotyl when *α1-COP* was absent **(Figure 7C, D)**. A recent study has shown that lipid raft compositions are critical for the secretory movement of GPI-anchored PdBG2 to PD (Iswanto et al., 2020).

**Figure 7.**
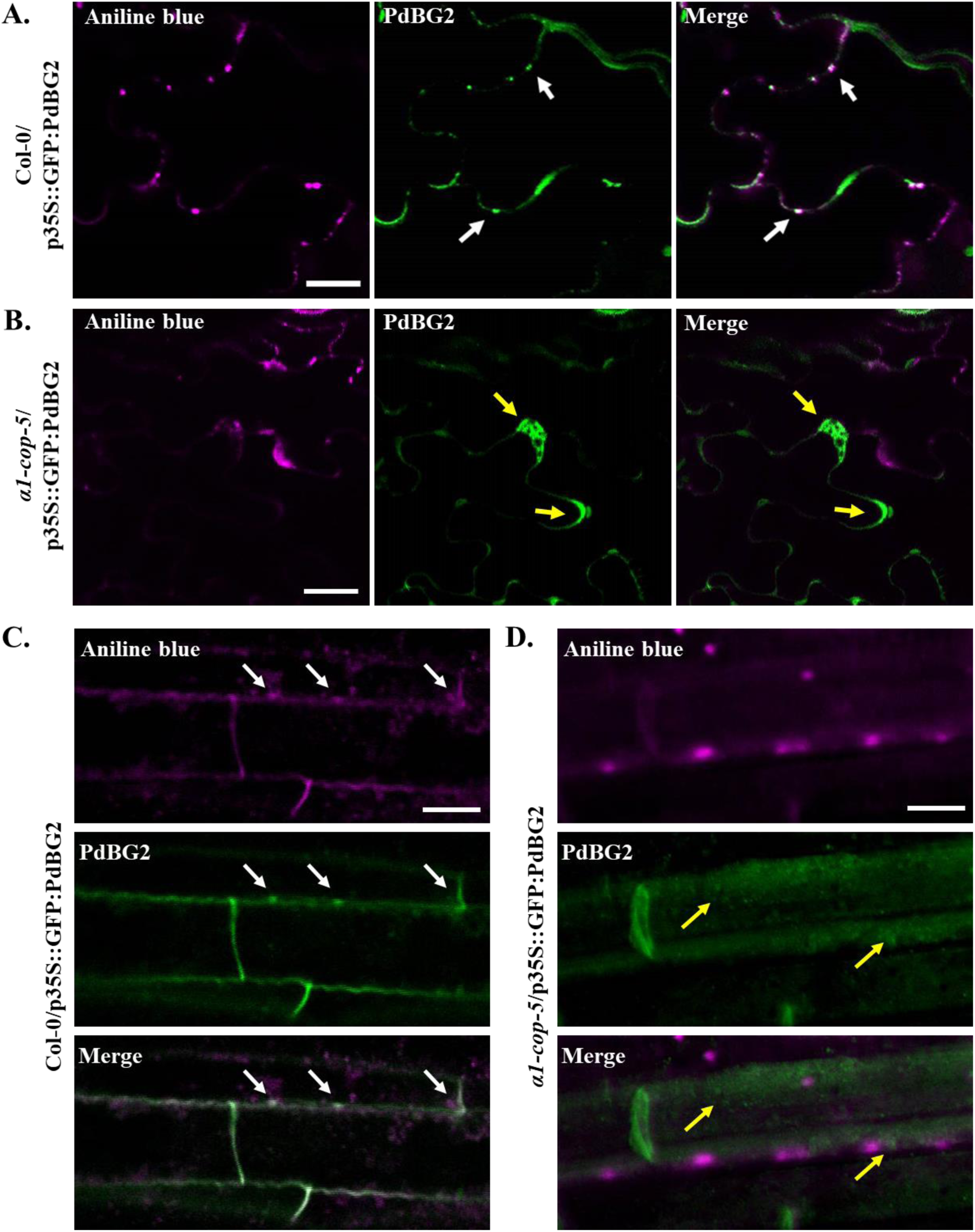
Subcellular localization of PdBG2 protein is altered in the *α1-cop-5* mutant. **(A, B)** Transgenic GFP:PdBG2 expression in the cotyledon of Arabidopsis wild-type Col-0 **(A)** and *α1-cop-5* mutant **(B)**. PdBG2 co-localizes with aniline blue-stained callose at PD in wild-type Col-0 (white arrows), but not in *α1-cop-5* mutant (yellow arrows). Scale bars: 20 µm. **(C, D)** Transgenic GFP:PdBG2 expression in the hypocotyl of Arabidopsis wild-type Col-0 **(C)** and *α1-cop-5* mutant **(D)**. PdBG2 co-localizes with aniline blue-stained callose at PD in wild-type Col-0 (white arrows), but not in *α1-cop-5* mutant (yellow arrows). Scale bars: 10 µm.

Next, the question arises whether α1-COP also plays a critical role in the translocation of non-GPI-anchored PD proteins. To answer the above question, the subcellular localization of PDLP1 and PDLP2 proteins in the absence of *α1-COP* was determined. Interestingly, subcellular localization of PDLPs proteins was not changed in the absence of *α1-COP* **(Supplemental Figure 9)**, indicating that α1-COP is not involved in the secretory cargo machinery of non-GPI-anchored PDLP1 or PDLP2 proteins. These data suggest that *α1-COP* is required for the recruitment of PdBG2 to PD. Moreover, the mislocalization of PdBG2 in the absence of α1-COP protein potentially led to the enhanced callose phenotype. An increase in callose level in the *α1-cop* mutants is not due to the decreased transcript of known PD specific callose degrading enzymes, namely, *BG_ppap*, *PdBG1*, *PdBG2*, and *PdBG3* **(Supplemental Figure 10)**. Taken together, current findings suggest that the α1-COP modulates the PD callose by regulating the secretion of PdBG2 to PD.

### *ERH1* and *GCS*-mediated PD callose regulation requires *α1-COP*

To gain further insight into the functions of *α1-COP, ERH1,* and *GCS* in callose-regulated symplastic continuity, PD callose was analyzed in plants overexpressing ERH1 and GCS, respectively. Interestingly, *ERH1* and *GCS* overexpression plants showed a significant reduction of callose depositions as compared with wild-type Col-0 **(Figure 8D-E)**. Furthermore, callose level in *erh1-1* (SALK_206784) and *gcs-2* mutants (Iswanto et al., 2020) was also quantified. **(Figure 8A-C)**. The callose level was significantly high in the *gcs-2* mutant, but the loss of function of *ERH1* did not show a significant difference in PD callose as compared to wild type **(Figure 8D-E).** This presumably reflects functional redundancy of the two homologs of *ERH1* found in Arabidopsis genome, AT2G29525 *ERH1-like1* (*ERHL1*) and AT3G54020 (*ERHL2*) (Wang et al., 2008).

**Figure 8.**
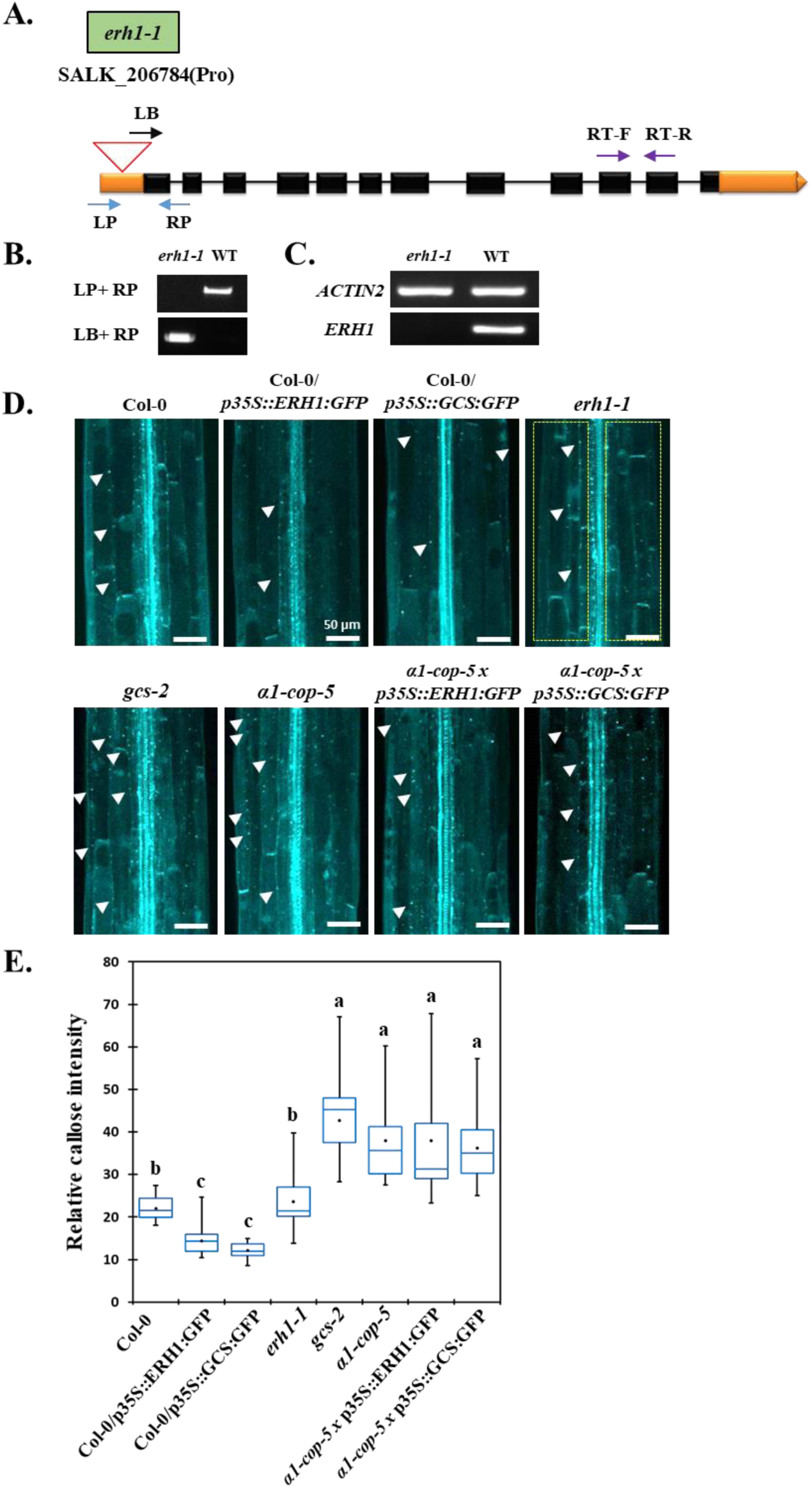
Excessive callose accumulation is maintained in the *α1-cop-5* overexpressing *ERH1* or *GCS* plants. **(A)** Gene structure of *ERH1* with one allele mutation. Two pair primers (LP-RP, and LB-RP) were used for T-DNA genotyping analysis. For RT-PCR analysis, one pair primer was designed in the exon parts of C-terminal region of *ERH1*. **(B)** T-DNA genotyping of *erh1-1* mutant using 2 pair primers (LP+RP) and (LB-RP), as shown in **(A)**. **(C)** RT-PCR analysis from wild type and *erh1-1* mutant. The amplicons were obtained from one pair primer as shown in figure **(A)**. *ACTIN2* was selected as reference gene. **(D)** Callose deposition analysis of Arabidopsis hypocotyls from each genotypes. Yellow square dotted lines were designated as region of interest (ROI) for measuring signal intensity. Scale bars: 50 µm. **(E)** Relative callose intensity quantification of Arabidopsis hypocotyls **(D)**. Statistical significance was done by One-Way ANOVA with Tuckey-Kramer test. (*n =* 25).

Since, the proper localizations of ERH1 and GCS require the biological function of α1-COP in Arabidopsis. We first asked if an attenuation in callose deposition actually requires the action of a α1-COP protein. For this study, crosses were made between *α1-cop-5* and Col-0/p35S::ERH1:GFP as well as *α1-cop-5* and Col-0/p35S::GCS:GFP. Homozygous F3 population from each genotypes were then examined for callose deposition analysis. Surprisingly, both *α1-cop-5/*p35S::ERH1:GFP and *α1-cop-5/*p35S::GCS:GFP plants showed significant increase in callose depositions in comparison to wild-type Col-0, *ERH1* and *GCS* overexpression plants. However, there were no significance differences in the observed callose depositions from *α1-cop-5, gcs-2, α1-cop-5/*p35S::ERH1:GFP and *α1-cop-5/*p35S::GCS:GFP plants **(Figure 8D-E)**. Collectively, these data suggest that α1-COP is essential for ERH1 and GCS functions in regulating callose deposition at PD.

### ERH1 and PdBG2 are the cargo proteins of α1-COP

In plants, lipid raft-enriched vesicle is required for GPI-anchored PdBG2 translocation (Iswanto et al., 2020). The secretory pathway of GPI-anchored PdBG2 and non GPI-anchored PDLP1 protein are segregated from ER to Golgi which explicate that there are at least two cargo machineries in the early secretory pathway, lipid raft and non lipid raft dependent manners (Iswanto and Kim, 2017; Iswanto et al., 2020). Moreover, non GPI-anchored PDLP1 protein interacts with α2-COP protein, not α1-COP protein (Caillaud et al., 2014) which is indicating that α2-COP protein may be involved in the secretory pathway of non GPI-anchored PD protein, especially for PDLP1. Since α1-COP function is required for cellular localization of of ERH1, GCS and PdBG2, thus we hypothesized that α1-COP is presumably associated with GPI-anchored PD proteins within intracellular compartment. To test the hypothesis, we examined if α1-COP is enriched by ERH1, then different combination of ERH1:GFP, α1-COP-RFP and VAMP721-RFP (vesicle marker) were transiently expressed in the *N. benthamiana* leaves with or without Exo1, an ER to Golgi intracellular trafficking blocker (Iswanto et al., 2020). Confocal imaging showed that when GFP:α1-COP was co-expressed together with VAMP721-RFP in the absence of Exo1, the fluorescent signals were partially co-localized at cell periphery **(Figure 9A)**, whereas in the presence of Exo1, the fuorescent signals were highly co-localized in the cytoplasm **(Figure 9B)**. Similarly, when ERH1:GFP was co-expressed together with VAMP721-RFP in the absence or presence of Exo1, the fluorescent signals were co-localized in the cell periphery or cytoplasm, respectively **(Figure 9C, D)**. Co-localization fluorescent signals were also observed when ERH1:GFP and α1-COP:RFP were co-expressed together with or without Exo1 **(Figure 9E, F)**, whereas the fluorescent signlas depicted from ERH1:GFP and α2-COP:RFP localization after Exo1 treatment did not show co-localization **(Supplemental Figure 11)**. Together, these results agree with BiFC and Co-IP results, validating that α1-COP physically interacts with ERH1 and specifically serves the cargo machinery of SL modifier ERH1.

**Figure 9.**
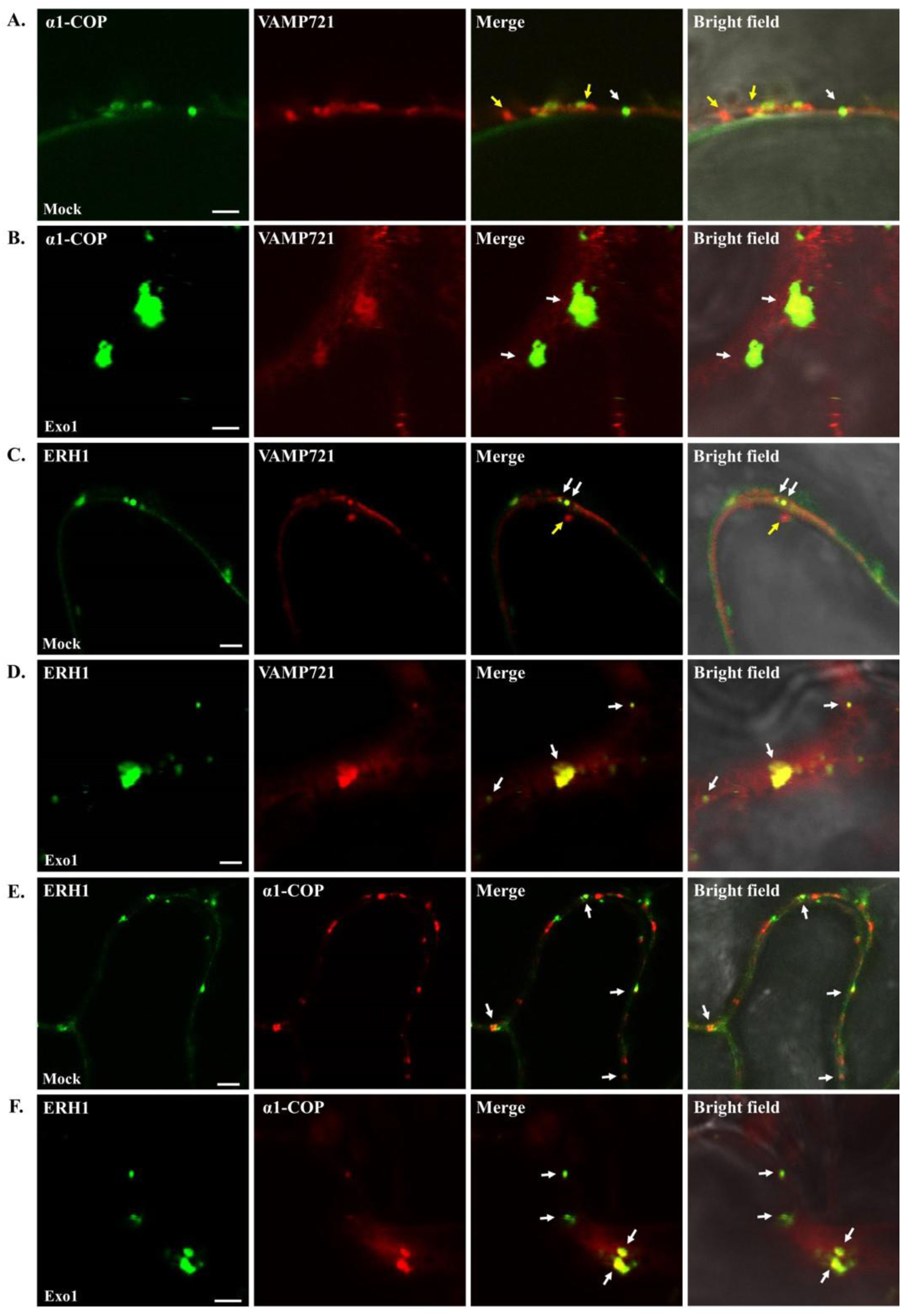
Subcellular localization of α1-COP and ERH1 in the presence of Exo1. **(A)** Confocal images of cells transiently expressing GFP:α1-COP and VAMP721:RFP in mock condition. **(B)** Confocal images of cells transiently expressing GFP:α1-COP and VAMP721:RFP in the presence of Exo1. **(C)** Confocal images of cells transiently expressing ERH1:GFP and VAMP721:RFP in mock condition. **(D)** Confocal images of cells transiently expressing ERH1:GFP and VAMP721:RFP in the presence of Exo1. **(E)** Confocal images of cells transiently expressing ERH1:GFP and α1-COP:RFP in mock condition. **(F)** Confocal images of cells transiently expressing ERH1:GFP and α1-COP:RFP in the presence of Exo1. *N. benthamiana* epidermal cells were used for colocalization studies. White arrows indicate colocalization of GFP and RFP signals, and yellow arrows indicate non colocalization. Scale bars **(A-F)** = 2 µm.

We next performed same experiments to determine whether PdBG2 is also the cargo protein for α1-COP. We co-expressed GFP:PdBG2 and α1-COP:RFP or PDLP1:GFP and α1-COP:RFP in the *N. benthamiana* leaves with or without Exo1. Confocal imaging exhibited that when GFP:PdBG2 or PDLP1:GFP was transiently co-expressed together with α1-COP:RFP in the absence of Exo1, the fluorescent signals were strongly co-localized at PD **(Figure 10A, C)**. Interestingly, the fluorescent signals depicted from GFP:PdBG2 and α1-COP-RFP were also co-localized at cytoplasm after Exo1 treatment, whereas PDLP:GFP signals were segregated from α1-COP:RFP signals in the presence of Exo1, not for GFP:PdBG2 **(Figure 10B, D)**. These results prompted us to deeply investigate the role of α1-COP or α2-COP in the protein cargo machinery of GPI-anchored PD protein and non GPI-achored PD protein. We next co-expressed GFP:PdBG2 and α2-COP:RFP or PDLP1:GFP and α2-COP:RFP in the *N. benthamiana* leaves with or without Exo1. Confocal imaging showed that when GFP:PdBG2 or PDLP1:GFP was co-expressed together with α2-COP:RFP in the absence of Exo1, the fluorescent signals were co-localized at PD **(Figure 10E, G)**. In contrast with α1-COP, we did not observe a co-localization of GFP:PdBG2 and α2-COP:RFP in the cytoplasm after Exo1 treatment. Surprisingly, after Exo1 treatment, PDLP1:GFP was highly co-localized with α2-COP:RFP in the cytoplasm **(Figure 10F, H)**. Collectively, our results suggest that α1-COP and α2-COP proteins independently involved in the proteins cargo machinery of GPI-anchored PdBG2 and non GPI-anchored PDLP1 proteins, which α1-COP is particularly subjected to the lipid raft-dependent manner **(Figure 12)**.

**Figure 10.**
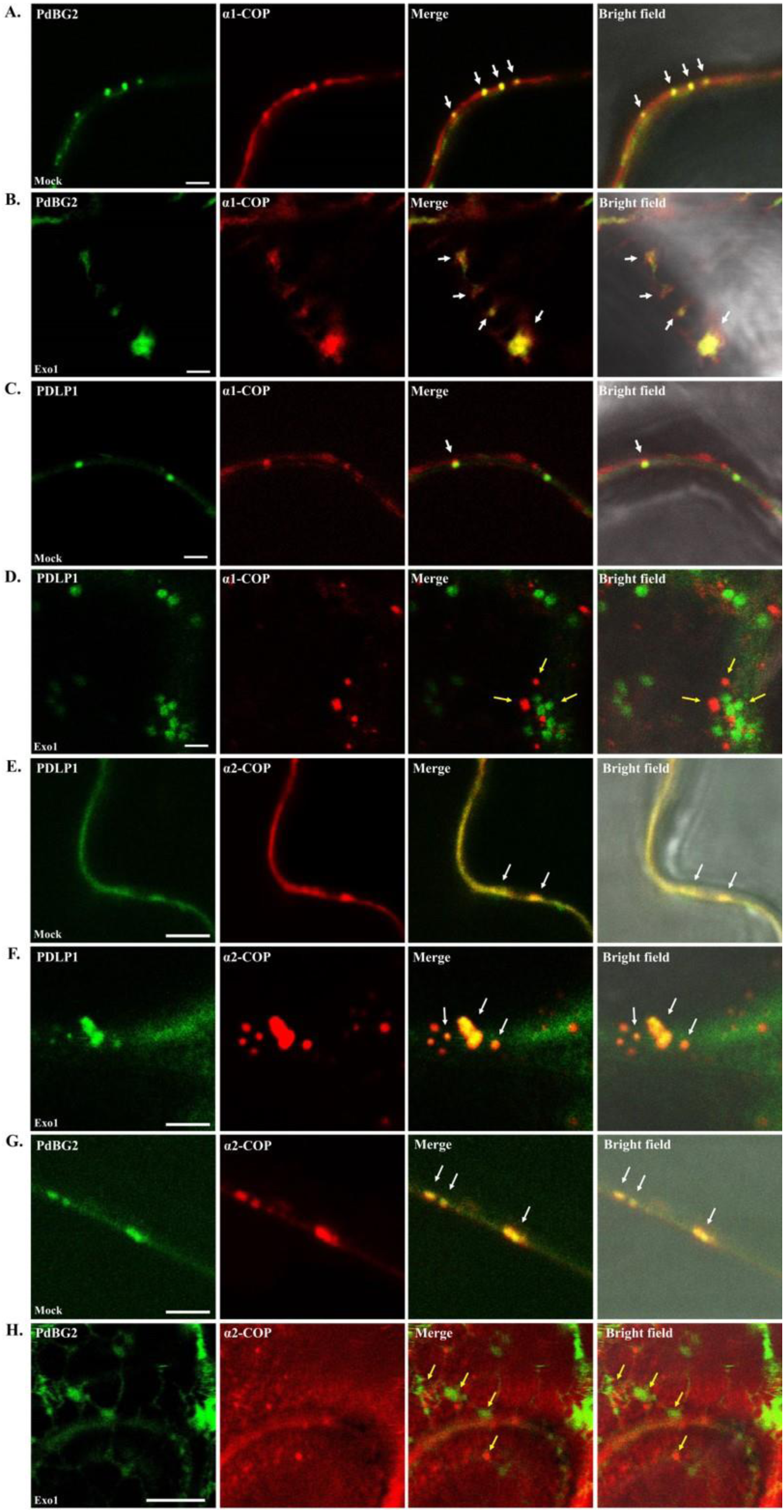
PdBG2 and α1-COP are retained in the same cellular compartment. **(A, B)** Confocal images of cells transiently expressing GFP:PdBG2 and α1-COP:RFP in mock condition **(A)** and Exo1 treated condition **(B)**. **(C, D)** Confocal images of cells transiently expressing PDLP1:GFP and α1-COP:RFP in mock condition **(C)** and Exo1 treated condition **(D)**. **(E, F)** Confocal images of cells transiently expressing PDLP1:GFP and α2-COP:RFP in mock condition **(E)** and Exo1 treated condition **(F)**. **(G, H)** Confocal images of cells transiently expressing GFP:PdBG2 and α2-COP:RFP in mock condition **(G)** and Exo1 treated condition **(H)**. *N. benthamiana* epidermal cells were used for colocalization studies. White arrows indicate colocalization of GFP and RFP signals, and yellow arrows indicate non colocalization. Scale bars = 2 µm in A-D, 5 µm in E-H.

### *α1-cop* mutants is susceptible against to *Botrytis cinerea*

Several studies have shown the link between SLs and callose homeostasis in response to biotic stimuli (Jacobs et al., 2003; Nishimura et al., 2003; Wang et al., 2008; Ellinger et al., 2013; Fang et al., 2016). We examined the possible involvement α1-COP activity in the plant defense response. First we grew and observed the intact phenotype of wild-type Col-0, *α1-cop* mutants along with *α1-COP* overexpression plants in normal condition, however we did not find a distinct phenotype, all genotypes were similar **(Supplemental Figure 12)**. Next we challenged the loss of function of *α1-COP* and overexpression plants with *B. cinerea* (a necrotrophic fungus). Leaves from *α1-cop* mutants showed severe lesion of fungus infection. The diameters area of necrotic lesion in *α1-cop-1* and *α1-cop-5* mutants were significantly larger than in wild-type Col-0 and *α1-COP* overexpression plants **(Figure 11A, B)**. In the previous study, a PD receptor-like protein, named LYSIN MOTIF DOMAIN-CONTAINING GLYCOSYLPHOSPHATIDYLINOSITOL-ANCHORED PROTEIN 2 (LYM2) is suggested to be an essential factor during *B. cinerea* infection in Arabidopsis (Faulkner et al., 2013; Vu et al., 2020). Since α1-COP is required for the proper intracellular trafficking of GPI-anchored PdBG2, we hypothesize that α1-COP is presumably associated with GPI-anchored LYM2 functioning. Thus, the enhancement of susceptibility of *α1-cop-1* and *α1-cop-5* plants against *B. cinerea*, may due to the malfunction of LYM2 protein. Taken together, this result indicates that the excessive callose deposition caused by the absence of α1-COP function results in a reduced plant defense response against to *B. cinerea*.

**Figure 11.**
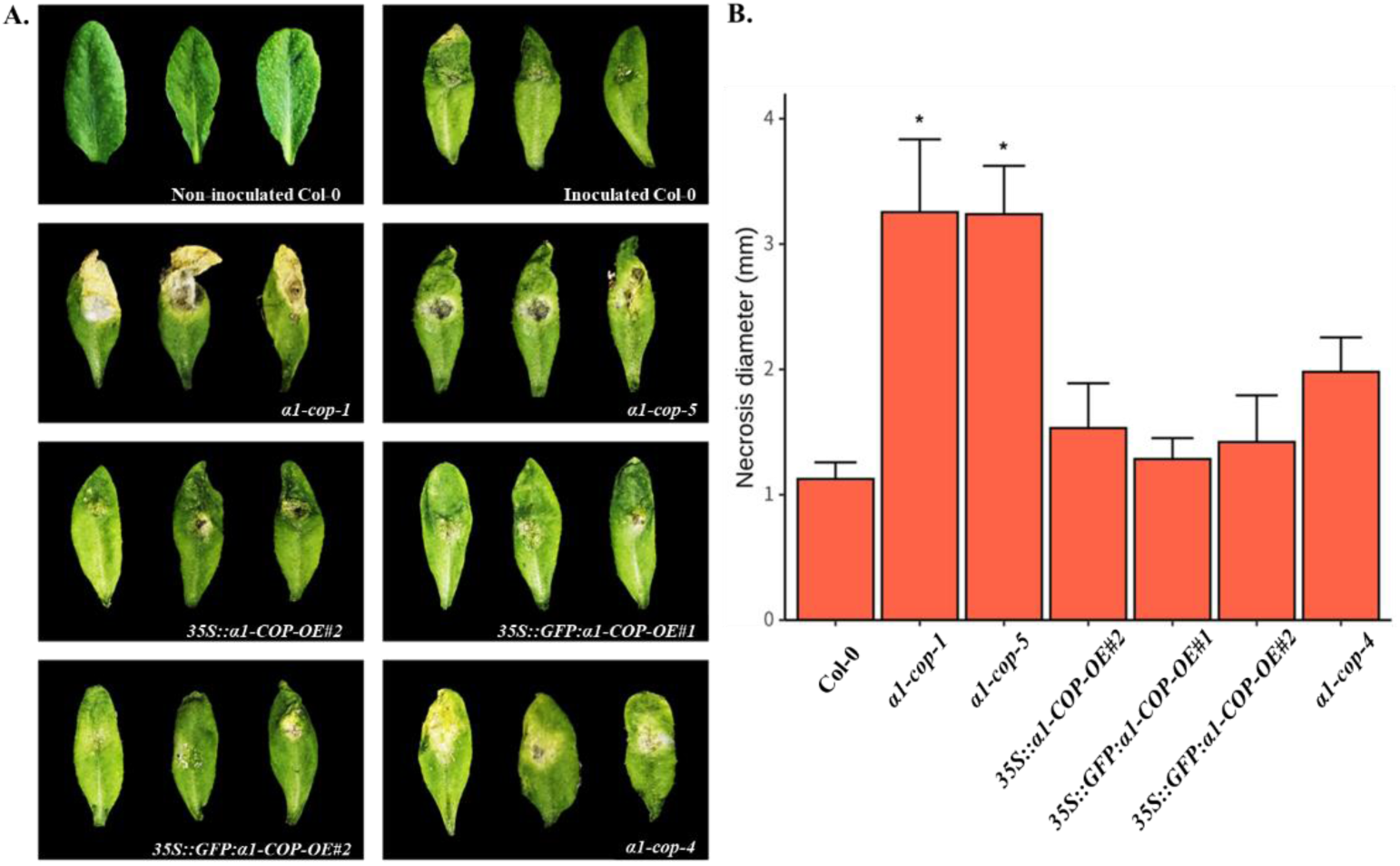
Susceptibility analysis of *α1-cop* mutants against to *Botrytis cinerea*. **(A)** Disease symptoms observation in 5 days post infection (dpi). **(B)** Leaf necrosis diameter were calculated at 5 dpi by measuring three leaves per plants for 5 plants. Two-sided Dunnett’s Multiple Comparisons was performed to determine significant difference with wild-type Col-0. (\**P* < 0.05).

## DISCUSSIONS

### Both α1-COP and α2-COP are partially located at PD, but only α1-COP is involved in callose-mediated phototropism

COPI is comprised by seven subunits (α/β/β’/γ/δ/ε/ζ) that have been classified into two sub complexes, the B- (α/β′/ε) and F-sub complex (β/δ/γ/ζ). In mammals, all the coatomer subunits have only one isoform, except γ-COP and ζ-COP subunits, whereas yeast contains only one isoform for all coatomer subunits. In contrast to mammals and yeast, in *A. thaliana* and other higher plants, except γ-COP and δ-COP subunits, every coatomer subunits have more than one isoform. Two α-COP isoforms, α1- and α2-COP, have been characterized in Arabidopsis (Gimeno-Ferrer et al., 2017; Cabada Gomez et al., 2020). Here we showed that a missense mutation (G486D) and knockout T-DNA mutants of *α1-COP* exhibited excess callose accumulation **(Fig. 1)** and had no or little defect in plant growth under normal condition **(Supplemental Figure 12)**. In contrast to *α1-cop* mutants, loss-of-function of *α2-COP* resembled wild-type Col-0 callose phenotype **(Supplemental Figure 3)**, but the plant growth is severely impaired (Gimeno-Ferrer et al., 2017). These two α-COP isoforms harbor the WD40 domain at their N-terminal, which is required for intracellular trafficking of cargo proteins containing KKXX motif. Interestingly, as these isoforms share an amino acid sequence of 93% identity, the excessive callose phenotype and the absence of growth defects in the *α1-cop* mutants might be explained by their differences at the subcellular localization, COPI composition and cargo specificity of both isoforms. In the subcellular localization study, we have shown that both α1-COP and α2-COP partially located at PD channels **(Figure 2, Supplemental Figure 13)**. Intriguingly, α2-COP is also detected in the nucleus **(Supplemental Figure 13)**, moreover, since α1-COP interacted with ε1-COP, β’2-COP and δ-COP, we strikingly found that α2-COP did not interact with β’2-COP, suggesting that α2-COP may has different COPI subunit to generate COPI complex **(Supplemental Figure 14A)**. Thus, PD callose phenotype should be resulted from other different nature between α1-COP and α2-COP. For their cargo specificity, two sphingolipid enzymes exposing KKXX dilysin motif on their C-terminal tail (ERH1 and GCS) have been shown to be α1-COP-dependent cargo machinery, but not α2-COP-dependent one **(Figure 12, Supplemental Figure 14B)**, suggesting that dilysin motif itself is not sufficient for COPI recruitment. Nonetheless, further studies will be required to identify the differences between the functions of *α1-COP* and *α2-COP* in the callose turnover and plant growth. The results presented here indicate that α1-COP has a role in callose-regulated symplasmic continuity.

**Figure 12.**
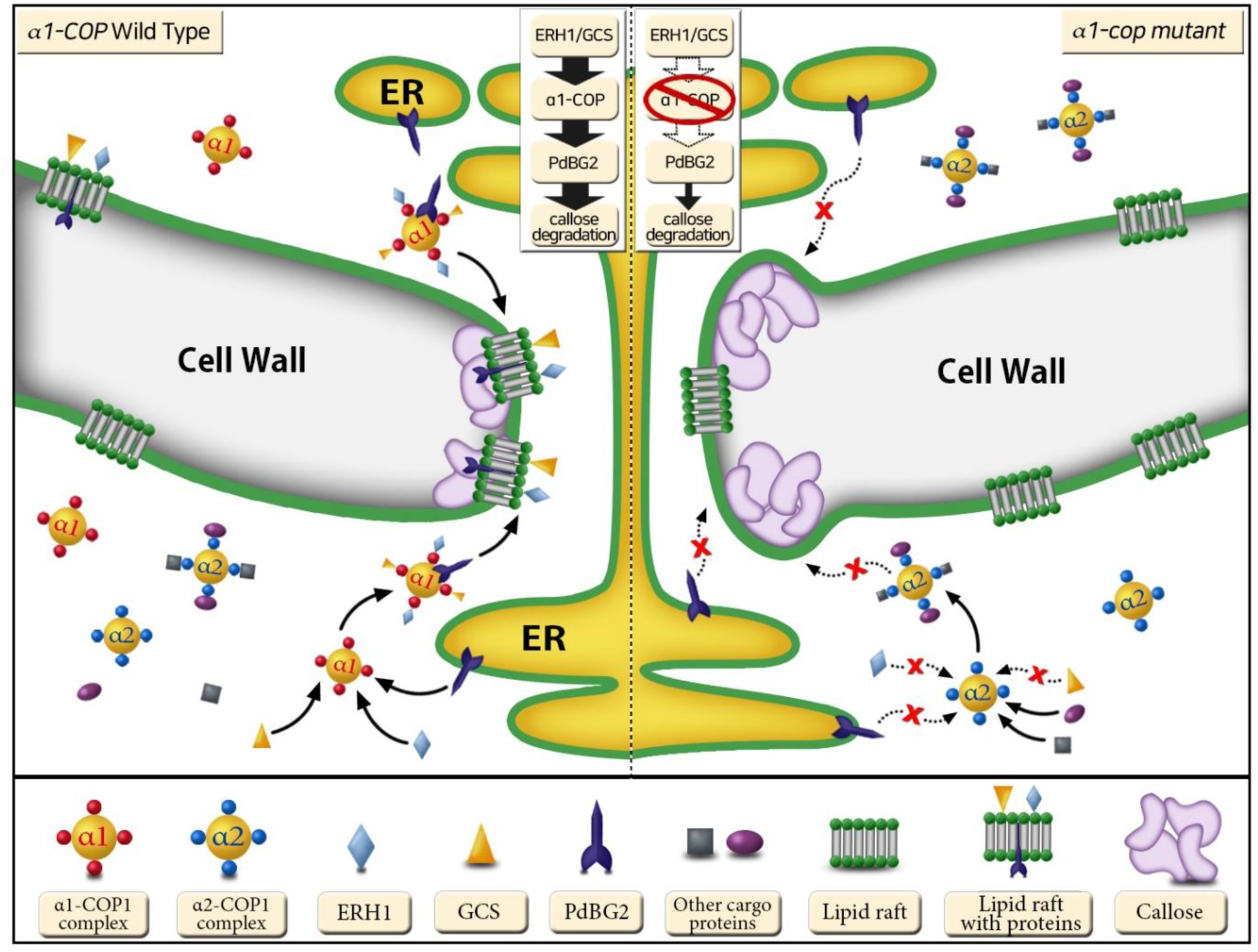
Schematic model of the role of α1-COP in the regulation of the PD. Sphingolipids are required for the formation of lipid rafts at PD-PM. SL modifier enzymes ERH1 and GCS facilitate the formation of more complex SLs-modulated lipid rafts. However, the translocation of ERH1 and GCS is dependent on an existence of α1-COP protein (not α2-COP protein) through direct binding activity. In the absence of α1-COP, ERH1 and GCS are not found at PD eventually alters lipid raft composition at PD-PM. The alteration of lipid rafts composition particularly effects on the subcellular localization of PdBG2 (callose degrading enzyme) leading a stabilization of callose deposition.

### α1-COP regulates intracellular trafficking of GPI-anchored PdBG2 through lipid raft dependent pathway

COPI vesicles are involved in several different intracellular transports of secretory proteins such as anterograde transport within the Golgi stack (Rothman, 1994; Orci et al., 1997), ER to Golgi transport (Pepperkok et al., 1993; Bednarek et al., 1995), and retrograde transports of GPI-anchored proteins (Sutterlin et al., 1997) and particular proteins that contain dilysine (KKXX) motif in their cytosolic C-terminal (Eugster et al., 2004; Jackson et al., 2012). In addition, it has been proven that COPI interacts with several sphingolipid molecules (Chaudhary et al., 1998; Contreras et al., 2012). Arabidopsis ERH1 and GCS proteins are two key SL pathway enzymes that are responsible in the conversion of ceramide species to produce inositolphosphorylceramide and glucosylceramide, respectively (Wang et al., 2008; Msanne et al., 2015). They contain dilysine motif at their C-terminus, which interact with COPI via α1-COP subunit **(Figure 3)**. ERH1 and GCS mainly localizes to the Golgi, ER as well as partially at PD. Here, we also found that loss of *α1-COP* causes obvious defects in trafficking of ERH1 and GCS proteins, which mostly localized to cytoplasm and were not found in the PD callose spot. This probably reflects the inability of ERH1 and GCS to enter into early secretory pathway under the absence of α1-COP. These results suggest the role of a specific *α1-COP* type of COPI in maintaining normal cellular function and intracellular trafficking of dilysine motif proteins, especially for several sphingolipid enzymes in Arabidopsis.

GIPCs and GlcCers are the most abundant sphingolipid species found in plasma membrane and other endomembranes which particularly required in the lipid raft formation. Moreover, PD-PM is enriched by lipid raft (Grison et al., 2015; Iswanto and Kim, 2017), and it have been reported that the proper form of lipid raft at PD is required for the GPI-anchored PD proteins localization such as PDCB1 and PdBG2 for maintaining callose level at PD neck region (Grison et al., 2015; Iswanto and Kim, 2017). Result from SLs analysis revealed that the reduction of GlcCers and GlcHCers molecules in the *α1-cop* mutants indicates the positive role of *α1-COP* in the SL biosynthesis pathways **(Figure 6)**. In yeast studies, it has been reported that mutations in the *ret1-1* (*α-cop*) disturb the ER to Golgi transport of GPI-anchored proteins (Sutterlin et al., 1997). Interestingly, we also found that loss of *α1-COP* changes the subcellular localization of GPI-anchored PdBG2 which was mostly detected in the cytoplasm and was absence at PD **(Figure 7)**. This probably reflects PdBG2’s inability to enter standard COPI vesicles for its ER to Golgi anterograde transport in *α1-cop* mutants. The absence of GPI-anchored PdBG2 at PD channel might be caused from the mislocalizations of ERH1 and GCS in particular to provide a biological property at PD-PM.

In summary, all these results suggest that α1-COP plays a role in targeting GPI-anchored protein to PD-PM and in modulating callose turnover through physical interaction with SL modifiers and their delivery in Arabidopsis. This work provides a key clue in our understanding of PD regulation by COPI vesicle functioning especially in the intracellular trafficking pathways.

## METHODS

### Plant materials and growth conditions

The mutants used in this study were in the *A. thaliana* wild-type Col-0 ecotype. The *α1-cop-4* (single amino acid substitution, Glycine to Aspartic acid, G486D) mutant was obtained from M4 population of *dsGSL8*-RNAi treated with ethyl methanesulfonate (EMS). The T-DNA insertion lines *α1-cop-1* (SALK_078465), *α1-cop-5* (SALK_003425), *α2-cop-1* (SALK_103968), *α2-cop-2* (SALK_1229034)*, erh1-1* (SALK_206784) and *gcs-2* (CS10111117) were obtained from the ABRC Arabidopsis stock center. These mutants were verified by PCR analysis using with T-DNA specific and flanking primers. The seeds were surface sterilized with 25% (v/v) bleach for 15 min, washed four times with sterile water, and kept in darkness at 4 °C for 3 days before they were planted on agar Murashige and Skoog (MS) medium. Plants were grown at 22 °C under 16 h light/8 h dark cycle.

### Plasmid Constructs

To create stable lines overexpressing *α1-COP*, *ERH1* and *GCS* or to perform transient localization assay of α1-COP, α1-COP^G486D^, ε1-COP, β’2-COP, δ-COP, ERH1, GCS, PDLP1, PDLP2 and PDLP5. PCR products (with/without stop codon) amplified from coding sequence (CDS) were first cloned in the pDONR207 plasmid (Invitrogen). The resultant entry clones were subsequently transformed into gateway binary vectors; pMDC43, pMDC83 (Curtis and Grossniklaus, 2003), pH7RWG2.0 (Karimi et al., 2002), myc-pBA, pDEST-^GW^VYNE, pDEST-VYNE(R), pDEST-^GW^VYCE and pDEST-VYCE(R)^GW^ (Gehl et al., 2009) to fuse GFP, RFP, Myc, Venus-N and Venus-C tags, respectively. PdBG2 construction was performed as previous described (Iswanto et al., 2020). To generate fusion protein of α2-COP, the introns-exons containing genomic DNA was amplified from ATG to (with/without) stop codon. PCR products were first cloned in the pDONR207 plasmid. The resultant entry clones were subsequently transformed into gateway binary vectors; pMDC43, pMDC83 and pDEST-VYCE(R)^GW^.

### Plant transformation and transgenic plant screening

Transgenic Arabidopsis plants were obtained by *Agrobacterium tumefaciens*-mediated transformation (Zhang et al., 2006). The developing Arabidopsis inflorescences were dipped 0.03% (v/v), 3% (m/v) sucrose and Agrobacterium cells carrying the chosen vectors for 5 seconds. T1 seeds were grown on selective media to screen transgenic Arabidopsis plants.

### *B. cinerea* infection assay

The plant was grown in a growth chamber under 12/12h light/dark at 22°C, 60% relative humidity. A challenging pathogen, *B. cinerea*, was cultivated on PDA (Potato Dextrose Broth 24 g, Agar 20 g, ddH_2_O 1 L) at 27°C for 7 days. Mycelium and spores were collected with sterile water and was filtered using the three layers of sterilized cheesecloth to collect spores. The spore concentration was adjusted to 1×10^5^ CFU/mL using the Hemocytometer (SUPERIOR, Lauda, DE). Rosette stage of *A. thaliana* of the wild type and the mutants were used in the experiment to confirm susceptibility against *B. cinerea*. The spore (5 uL; 10^5^ CFU/mL) was inoculated per leave of *A. thaliana*. The plant was covered with parafilm to avoid dispersion of conidia spore and to maintain high humidity (95-100%). Experimental repetition per line was performed three leaves per plants for 5 plants. After 5 days inoculation, necrosis symptoms were evaluated. Diameter on the leaves was measured using Image J program. Two-sided Dunnett’s Multiple Comparsions was performed to determine significant difference in disease incidence (*P* < *0.05*). Statistix 8 (version 8.0) was used as the analytical software.

### Aniline Blue Staining

Arabidopsis hypocotyls, root tips and rosette leaves were kept in callose staining buffer (CSB) for 3 h in darkness. CSB was a mixture of 0.1% (w/v) aniline blue in autoclaved triple-distilled water and 1 M glycine (pH 9.5) at a volume ratio of 2:3. The samples were then washed, and the fluorescence was detected under a confocal microscope. During quantification, yellow square dotted lines were selected as a region of interest (ROI) and the mean relative fluorescence intensity was measured using ImageJ (https://imagej.nih.gov/ij/). For additional information on quantification of callose using aniline blue staining, see (Zavaliev et al., 2011; Zavaliev et al., 2013).

### Hypocotyl Loading Assay

To measure symplasmic connectivity using the HPTS (8-Hydroxypyrene-1, 3, 6-trisulfonic acid trisodium salt, SIGMA-ALDRICH) dye movement assay, a symplasmic dye tracer, was loaded on the top of sharply trimmed etiolated three-day-old Arabidopsis hypocotyls as shown in previous publications (Han et al., 2014; Kumar et al., 2016). A cover slip was placed between each cut hypocotyl surface and the MS agar. For dye loading, individual agar blocks containing HPTS (5 mg/mL) were placed on the cut hypocotyl surface. After a 5 min loading period, the seedlings were washed in water for 15 min, and then fluorescent probe movements were observed by confocal microscopy (Kumar et al., 2016).

### RNA extraction and QRT-PCR analyses

Total RNA was extracted from 10-day-old Arabidopsis seedlings with an RNeasy^®^ Plant Mini Kit (QIAGEN) according to the manufacturer’s instructions. First-strand complementary DNA synthesis was performed using 1 µg of total RNA with an anchored oligo (dT) and Transcriptor Reverse Transcriptase (QIAGEN) following the manufacturer’s protocol. Quantitative RT-PCR was conducted on 384-well plates using the Light Cycler 480 system (Biorad) and the QuantiSpeed SYBR Green Kit (PhileKorea) under the following conditions: denaturation for 5 min at 95 °C, 40 cycles of 10 s at 95 °C for denaturation and 10 s at 60 °C for annealing. Each reaction was performed with 3 µL of 1:20 (v/v) dilution of the first complementary DNA strand, with 0.5 µM of each primers **(Supplemental Table 1)** in a total reaction volume of 10 µL. The QRT-PCR data represent mean value of two independent biological experiments, with four technical replicates after normalization with the four reference transcripts (*ACTIN2*, *UBQ10*, *UBC9* and *EF-1α*) shown before to exhibit invariable expression levels.

### Bimolecular fluorescence complementation (BiFC) assays

The CDSs of *α1-COP, α2-COP, β’2-COP, ε1-COP, ε2-COP, δ-COP, ERH1, GCS* and *LOH1* were cloned into a set of binary BiFC-Gateway vectors; pDEST-^GW^VYNE (Venus aa 1-173), pDEST-VYNE(R)^GW^ (Venus aa 1-173), pDEST-^GW^VYCE (Venus aa 156-239) and pDEST-VYCE(R)^GW^ (Venus aa 156-239) with kanamycin selection marker in *E.coli* and *A. tumefaciens*. The combination of proteins (in the relevant figure) was transiently co-expressed in *N. benthamiana*. Venus fluorescent signals were observed at 72 h post infiltration under OLYMPUS FV1000-LDPSU (Olympus, Japan) confocal laser-scanning microscope.

### Co-immunoprecipitation (Co-IP) assay

The combination of proteins (in the relevant figure) were transiently expressed in *N. benthamiana*. For bead preparation, 75 µl of protein A agarose was washed in 500 µl IP buffer (100 mM Tris-HCl pH 7.5, 150 mM NaCl, 1 mM EDTA pH 8.0, 0.5% NP40, ddH_2_O, 3mM DTT) with 1:100 complete protease inhibitor cocktail (Roche) and centrifuged at 3000 rpm for 1 min at 4°C. The washing step was repeated for 3 times. The protein A agarose bead was added with 450 µl of IP buffer and 5 µl of Myc antibody (Cell Signaling). The mixture was then incubated on rotator at 4°C for 4 h. Next, 1 g fresh weight of infiltrated leaves were collected 3 days after infiltration. The tissues were ground in 3 ml IP buffer. The broken tissues were transferred to the filter and aliquoted into eppendorf tubes. The samples were centrifuged at 12.000 rpm for 5 min at 4°C to remove cell debris. 15 µl of supernatants (total proteins) served as the input controls. 450 µl of supernatants were then incubated with 55 µl of protein A agarose conjugated with Myc antibody on rotator at 4°C for 12 h. The protein A agarose beads were then spun down at 3000 rpm for 2 min at 4°C and washed three times with IP buffer. After the last centrifugation, supernatants were removed and beads were adjusted up to 40 µl with IP buffer. Proteins associated with the Myc-fusion proteins were eluted by adding 10 µl of 5x sample loading buffer and heating at 70°C for 5 min. The eluted proteins were analyzed by immunoblot assay. Primary antibodies used in this study were Myc-Tag (9B11) Mouse mAb (Cell Signaling) and Anti-GFP (abcam ab6556). Secondary antibodies used in this study were Anti-mouse IgG, HRP-linked antibody (Cell Signaling) and Anti-Rabbit IgG (H+L) Conjugate (Promega).

### Confocal Microscopy

Confocal fluorescence microscopy was performed with an OLYMPUS FV1000-LDPSU (Olympus, Japan) inverted confocal microscope using 20X/0.8 oil-immersion objective or 40X/1.3 oil-immersion objective. GFP was excited with a laser using 488 nanometer beam splitter. RFP and FM4-64 were excited with a laser using 543 nanometer beam splitter. YFP was excited with a laser using 515 nanometer beam splitter. Aniline blue and DAPI were excited with a laser using 405 nanometer beam splitter. Signal intensities from GFP, YFP and aniline blue (callose detection) were quantified with ImageJ software for statistical analyses.

### Sphingolipid extraction and sphingolipid analysis

Plant samples preparation for SL inhibitors treatments; Arabidopsis seeds from each genotypes; wild-type Col-0, *α1-cop-1*, *α1-cop-5*, α1-COP-OE#2 and α1-COP-OE#3 were grown on normal MS medium for 14 days. Two-week-old seedlings were immediately frozen in liquid nitrogen and ground to a fine powder (*n* = 250, 4 independent biological experiments). For plant sphingolipid analysis, the total lipids were extracted from 3 mg of lyophilized Arabidopsis seedlings using the combined upper phase (220 μL) and lower phase (110 μL) of methyl-*tert*-butyl ether (MTBE)/methanol/water (100:30:35, *v/v/v*) described previously (Chen et al., 2013). Extracts were reconstituted in 100 μL chloroform/methanol (1:9, *v/v*). Sphingolipids profiling was performed using a Nexera2 LC system (Shimadzu Corporation, Kyoto, Japan) connected to a triple quadrupole mass spectrometer (LC-MS 8040; Shimadzu, Kyoto, Japan) with reversed phase Kinetex C18 column (100 × 2.1 mm, 2.6 μm, Phenomenex, Torrance, CA, USA) for chromatographic separations of lipids. Mobile phase A consisted of water/methanol (1:9, *v/v*) containing 10 mM ammonium acetate, and mobile phase B consisted of isopropanol/methanol (5:5, *v/v*) containing 10 mM ammonium acetate. To achieve chromatographic separation, a gradient elution program was optimized as follows: 0 min, 30% B; 0-15 min, 95% B; 15-20 min, 95% B; 20-25 min, 30% B. The flow rate was set a 200 μL min^-1^. 5 μL sample volumes were injected for each run. To achieve sphingolipid quantifications, the calculated ratio of analyte and internal standard is multiplied by the concentration of the internal standard to obtain the concentration for each lipid species (Xia and Jemal, 2009; Bure et al., 2013; Lee et al., 2017; Im et al., 2019). Since there is no commercial internal standard for the quantification of GIPC molecular species, ganglioside GM_1_ is often used as an alternative to it (Markham and Jaworski, 2007; Tellier et al., 2014). We performed quantitative analysis of SLs using one-point calibrations of each target SL species [dihydrosphingosine d17:0/LCBs, non-hydroxy-phytoceramide (t18:0/8:0)/Ceramide, alpha-hydroxy-phytoceramide (t18:0/h6:0)/hydroxyceramide, glucosyl-ceramide (d18:1/12:0)/GlcCer or GlcHCer, and GM_1_ (d18:1/18:0)/GIPC with known concentration]. Non-hydroxy-phytoceramide [(t18:0/8:0), MW = 443.6] and alpha-hydroxy-phytoceramide [(t18:0/h6:0), MW = 431.6] were synthesized by Kyungpook University (Daegu, Korea) and other internal standards were purchased from Matreya (Pleasant Gap, PA, USA) or Avanti Polar Lipids (Alabaster, AL, USA).

### Data analyses and experimental repeats

The statistical analysis and sample size or number for each experiment were listed in the relevant figures and figure legends. All experiments in this study, at least were conducted with three independent replications, see **(Supplemental Table 2)** for the summary of statistical tests.

### Accession numbers

Accession numbers for the genes characterized in this work, see **(Supplemental Table 3)**.

## Supplemental Data

**Supplemental Figure 1.** Next Generation Mapping (NGM) and phototropic response analysis in the *α1-cop* mutants.

**Supplemental Figure 2.** α1-COP overexpression lines reduce callose level.

**Supplemental Figure 3.** Callose deposition and phototropic response analyses in the *α2-cop* mutants.

**Supplemental Figure 4.** α1-COP interacts with F-subcomplex and B-subcomplex members.

**Supplemental Figure 5.** Single amino acid substitution does not alter the subcellular localization of α1-COP.

**Supplemental Figure 6.** Arabidopsis wild-type Col-0 and *α1-cop-5* plants expressing ERH1:GFP.

**Supplemental Figure 7.** BFA disrupts intracellular transport of α1-COP and ERH1 proteins.

**Supplemental Figure 8.** Quantitative RT-PCR analyses of sphingolipid enzymes in the wild-type Col-0, *α1-cop-5* mutant and α1-COP-OE#2 overexpression plants.

**Supplemental Figure 9.** Subcellular localization of PDLP(s) proteins in the *α1-cop-5* mutant.

**Supplemental Figure 10.** Quantitative RT-PCR analyses of callose degradation enzymes in wild-type Col-0, *α1-cop-5* mutant and α1-COP-OE#2 plants.

**Supplemental Figure 11.** Subcellular localization of α2-COP and ERH1 in the presence of Exo1.

**Supplemental Figure 12.** Growth of wild-type Col-0, *α1-cop* mutants and α1-COP overexpression plants.

**Supplemental Figure 13.** Subcellular localization of α2-COP protein.

**Supplemental Figure 14.** The interaction partner analysis of α2-COP protein with COPI sub-complex members and sphingolipid modifier enzymes.

**Supplemental Table 1.** Primers used in this study.

**Supplemental Table 2.** Summary of statistical tests.

**Supplemental Table 3.** Accession numbers for the genes characterized in this work

## ACKNOWLEDGMENTS

This work was supported by the National Research Foundation of Korea Grant NRF 2018R1A2A1A05077295, 2020M3A9I4038352 and 2020R1A6A1A03044344.

## AUTHOR CONTRIBUTIONS

ABBI, MHV, RK, JCS, SW, DRK, KYS, SGH, HK, WYK, SHK, KHL and JYK conceived the study. ABBI. performed experiments, analyzed data and wrote the manuscript. ABBI. and JYK designed experiments. MHV, RK, JCS, SW, DRK, KYS, SGH, HK, WYK, SHK, and KHL performed experiments. All authors contributed to and edited the final manuscript.

## Notes

### Competing Interest Statement

The authors have declared no competing interest.

