## Supplemental Data for "α1-COP delivers sphingolipid modifiers and controls plasmodesmal callose deposition in Arabidopsis"

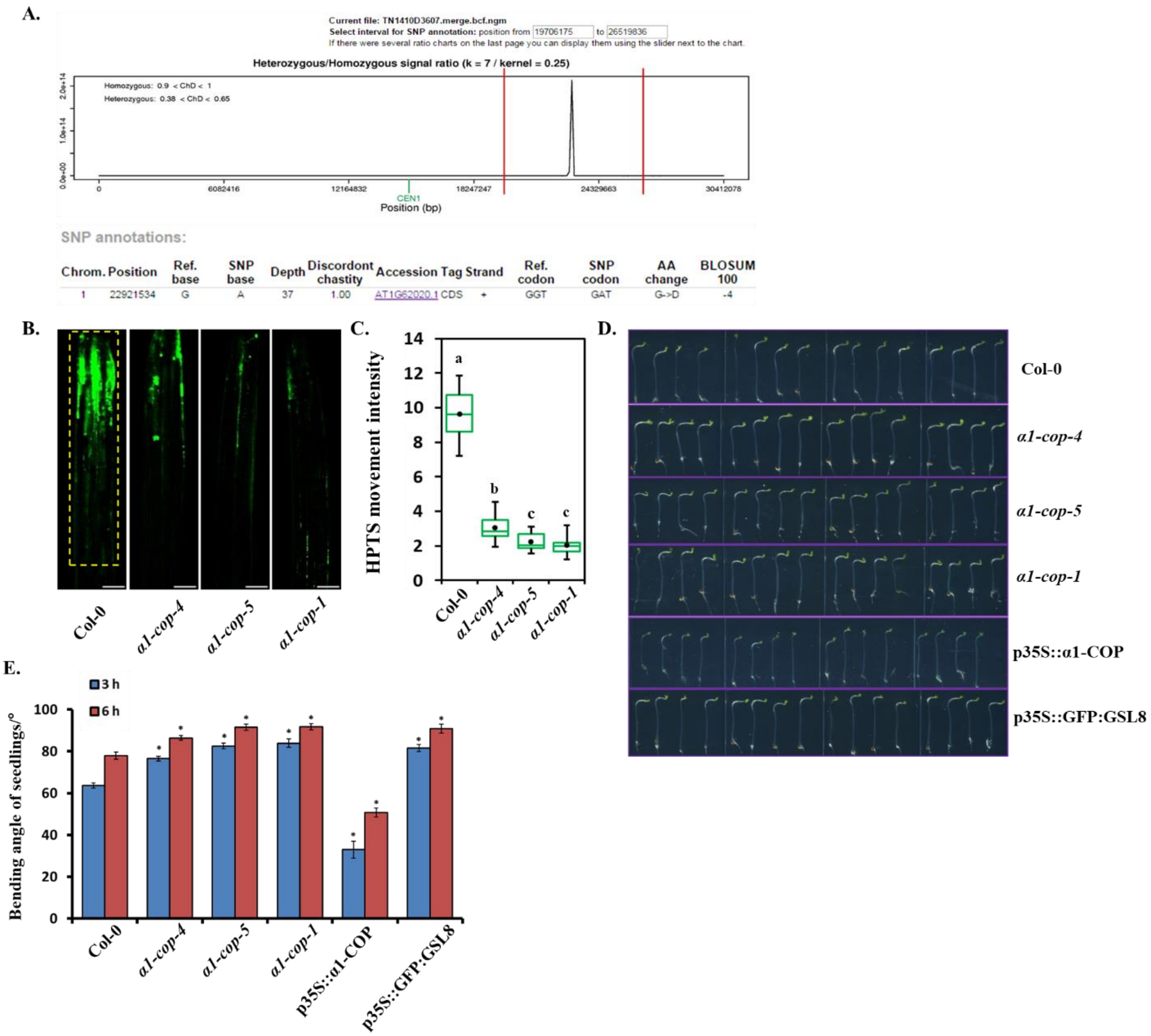


**Supplemental Figure 1. Next Generation Mapping (NGM) and phototropic response analysis in the *α1-cop* mutants.**

**(A)** NGM result showed that single amino acid from α1-COP was changed (Glycine (G) to Aspartic acid (D)).

**(B)** PD permeability analysis in the Arabidopsis hypocotyls. Three-day-etiolated seedlings of wild-type Col-0 and *α1-cop* mutants were analyzed using HPTS loading assay. Scale bars: 100 µm.

**(C)** HPTS fluorescence intensity quantification of Arabidopsis hypocotyls from wild-type Col-0, *α1-cop-4, α1-cop-5* and *α1-cop-1* plants. (*n*=10).

**(D)** Phototropic responses were performed by wild-type Col-0, *α1-cop* mutants, *α1-COP* and *GSL8* overexpression plants. Image was captured after 6 h of unilateral white-light illumination (2 μmol $m^{-2}$ $s^{-1}$).

**(E)** Phototropic hypocotyl bending angle measurement. Hypocotyl bending angles were measured under unilateral white-light illumination and the average bending angles from three independent experiments were calculated (n=16). Error bars represent standard error (SE). (Student's *t*-test between wild-type Col-0, *α1-cop* mutants, *α1-COP* and *GSL8* overexpression plants hypocotyls after 3 h and 6 h of unilateral light treatment, $^{*}P<0.05$).

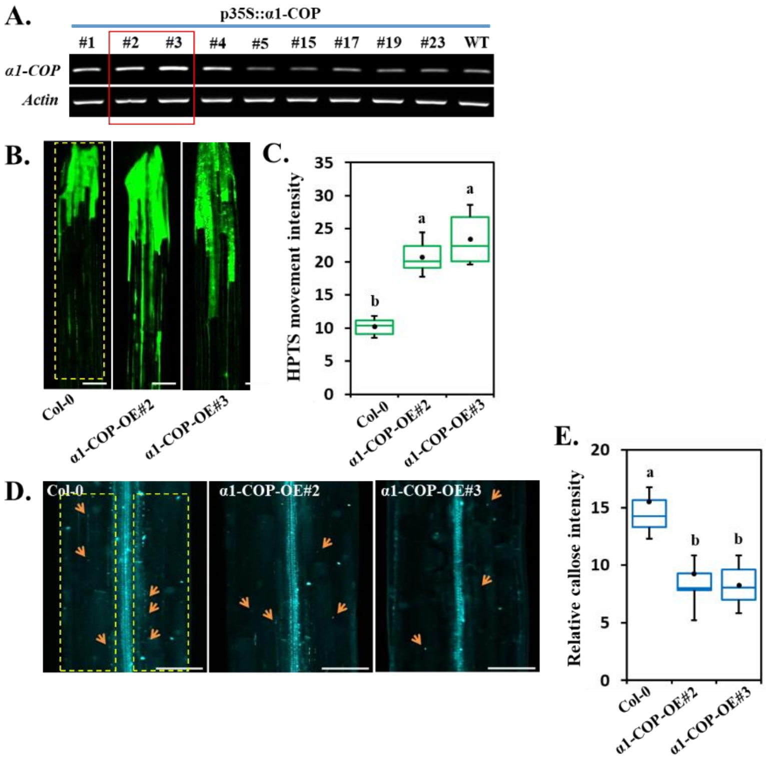


**Supplemental Figure 2. α1-COP overexpression lines reduce callose level.**

**(A)** RT-PCR analysis of p35S::α1-COP overexpression transgenic plants. One pair primers that were previously designed for checking *α1-COP* mRNA level in the mutant lines was also used for α1-COP overexpression lines screening. We selected α1-COP lines number #2 and #3 for further experiments.

**(B)** PD permeability assessment in the Arabidopsis hypocotyls of wild-type Col-0 and α1-COP overexpression transgenic plants (α1-COP-OE#2 and α1-COP-OE#3) using HPTS loading assay. Scale bars: 100 μm.

**(C)** Quantification of HPTS fluorescence intensity of wild-type Col-0 and α1-COP overexpression plants. (*n*=10).

**(D)** Callose deposition analysis of Arabidopsis hypocotyls of wild-type Col-0 and α1-COP overexpression plants using aniline blue staining. Scale bars: 100 μm.

**(E)** Quantification of relative callose intensity of wild-type Col-0 and α1-COP overexpression plants. Statistical significances **(C and E)** were done using One-Way ANOVA with Tuckey-Kramer test. (*n*=10). Yellow square dotted lines **(B and D)** were designated as region of interest (ROI) for measuring signal intensity.

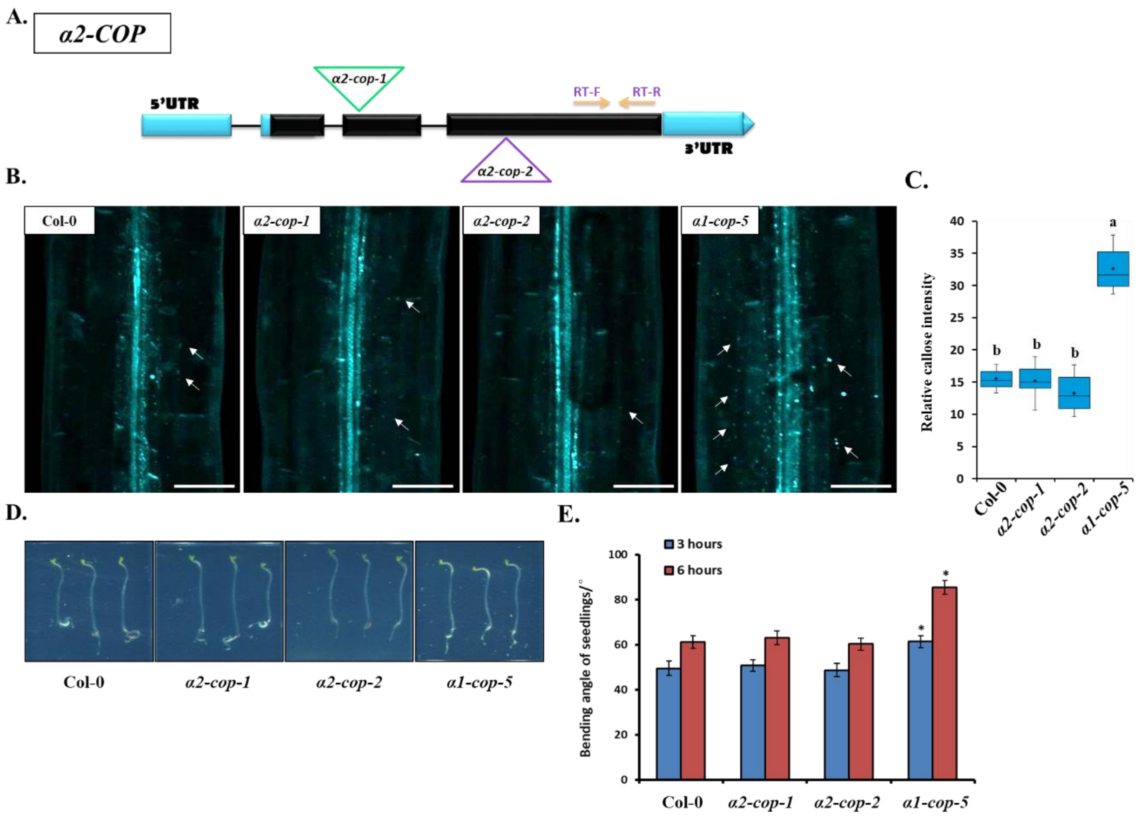


**Supplemental Figure 3. Callose deposition and phototropic response analyses in the α2-cop mutants**.

**(A)** Gene structure of *α2-COP* with two allele mutations. *α2-cop-1* and *α2-cop-2* were collected from ABRC as T-DNA insertion lines.

**(B)** Callose deposition analysis of Arabidopsis hypocotyls from wild-type Col-0, *α2-cop-1*, *α2-cop-2* and *α1-cop-5* mutants. Scale bars: 100 µm.

**(C)** Relative callose intensity quantification of Arabidopsis hypocotyls from wild-type Col-0, *α2-cop-1*, *α2-cop-2* and *α1-cop-5* mutants. (n=10).

**(D)** Phototropic responses were performed by wild-type Col-0, *α2-cop-1*, *α2-cop-2* and *α1-cop-5* mutants. Image was captured after 6 h of unilateral white-light illumination (2 µmol m−2 s−1).

**(E)** Phototropic hypocotyl bending angle measurement. Hypocotyl bending angles were measured under unilateral white-light illumination and the average bending angles from three independent experiments were calculated (n=16). Error bars represent standard error (SE). (Student's t-test between wild-type and *α1-cop-5* mutant hypocotyls after 3 h and 6 h of unilateral light treatment, *$P<0.05$).

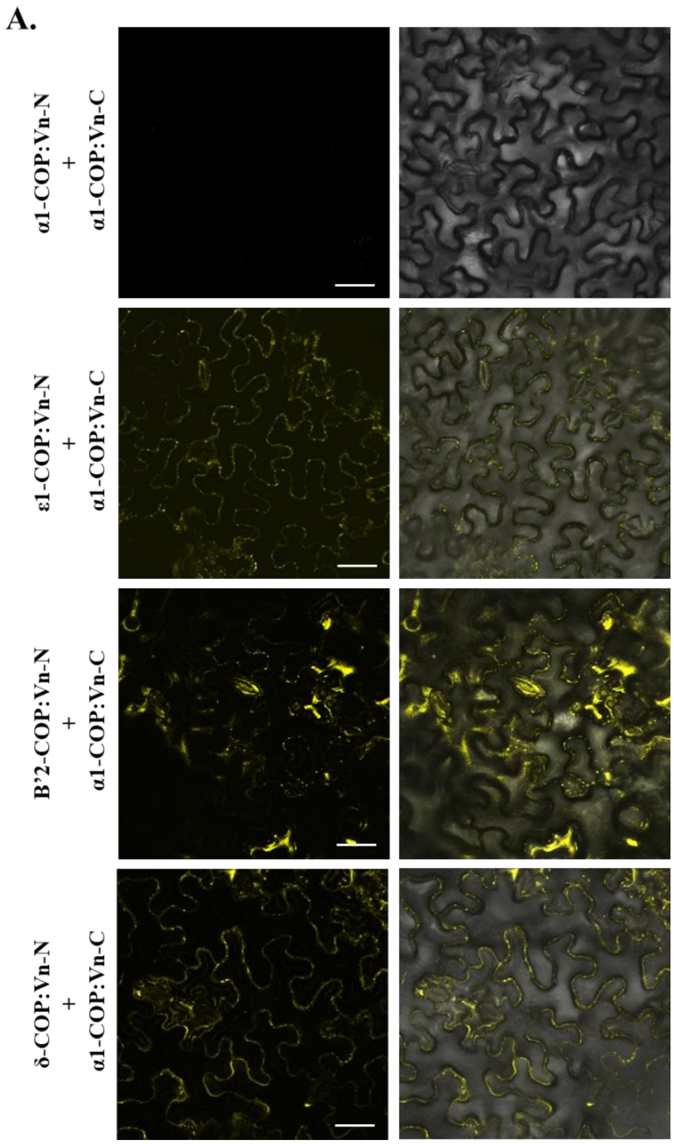


**Supplemental Figure 4. α1-COP interacts with F-subcomplex and B-subcomplex members.** **(A)** BiFC analysis of ε1-COP, β'2-COP, δ-COP interactions with α1-COP in *planta*. Fluorescence was observed in the transformed *N. benthamiana* leaf epidermal cells, which results from complementation of the N-terminal part of the Venus fused with ε1-COP, β'2-COP and δ-COP by the C-terminal part of the Venus fused with α1-COP. Scale bars: 50 µm.

A.

| Gene | Gene Identifier | Retrieval target motif (C-terminal domain) KKXX | Position |
|---|---|---|---|
| *LOH1* | AT3G25440 | KKKR | 441-444 |
| *LOH2* | AT3G19260 | - | - |
| *LOH3* | AT1G13580 | - | - |
| *ERH1* | AT2G37940 | KKIY, KKLP | 104-107, 242-245 |
| *ERHL1* | AT2G29525 | KKIC | 104-107 |
| *ERHL2* | AT3G54020 | KKIY, KKLP | 104-107, 243-246 |
| *GCS* | AT2G19880 | KKGA, KKFD | 492-495, 516-519 |

B.

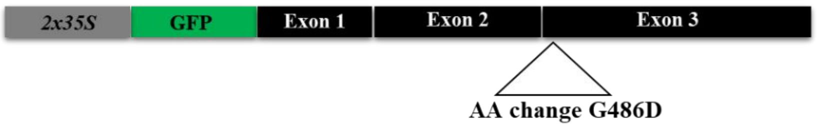


C.

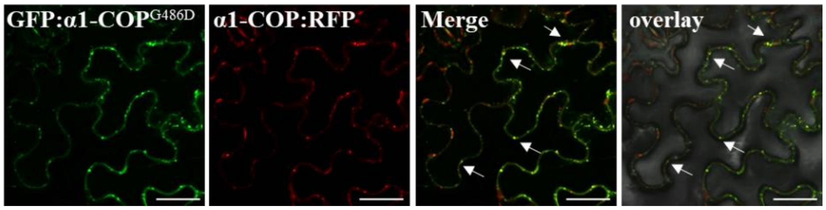


**Supplemental Figure 5. Single amino acid substitution does not alter the subcellular localization of α1-COP.**

**(A)** Table of retrieval target motif for several sphingolipid enzymes existing in *Arabidopsis thaliana*.

**(B)** Construct for the expression of α1- COP$^{G486D}$ fusion protein. GFP-tagged α1-COP$^{G486D}$ was generated by using Gateway cloning method.

**(C)** Confocal images of *N. benthamiana* epidermal cells transiently expressing fluorescent fusion proteins after agroinfiltration. Colocalization of GFP-α1-COP$^{G486D}$ and α1-COP-RFP. White arrows indicate co-localization of GFP and RFP signals. Scale bars: 50 µm.

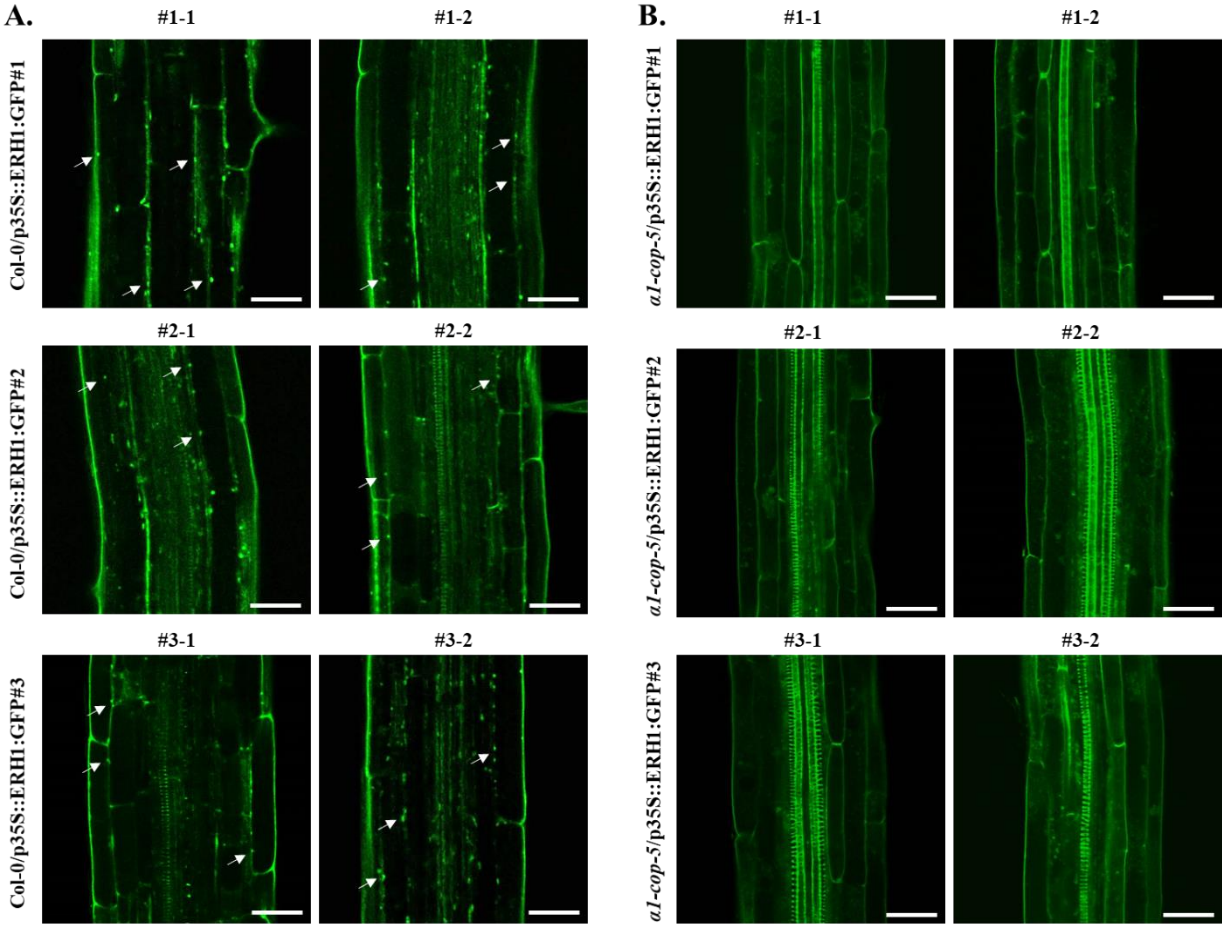


**Supplemental Figure 6. Arabidopsis wild-type Col-0 and *α1-cop-5* plants expressing ERH1:GFP.**

**(A)** Confocal images of T3 homozygous Col-0/ERH1:GFP lines #1, #2 and #3. GFP fluorescence signals were observed in the primary root of 9-day-old Arabidopsis seedlings. White arrows indicate the GFP signals in specific punctate spots. Scale bars: 30 µm.

**(B)** Confocal images of T3 homozygous *α1-cop-5*/ERH1:GFP lines #1, #2 and #3. GFP fluorescence signals were observed in the primary root of 9-day-old Arabidopsis seedlings. Scale bars: 30 µm.

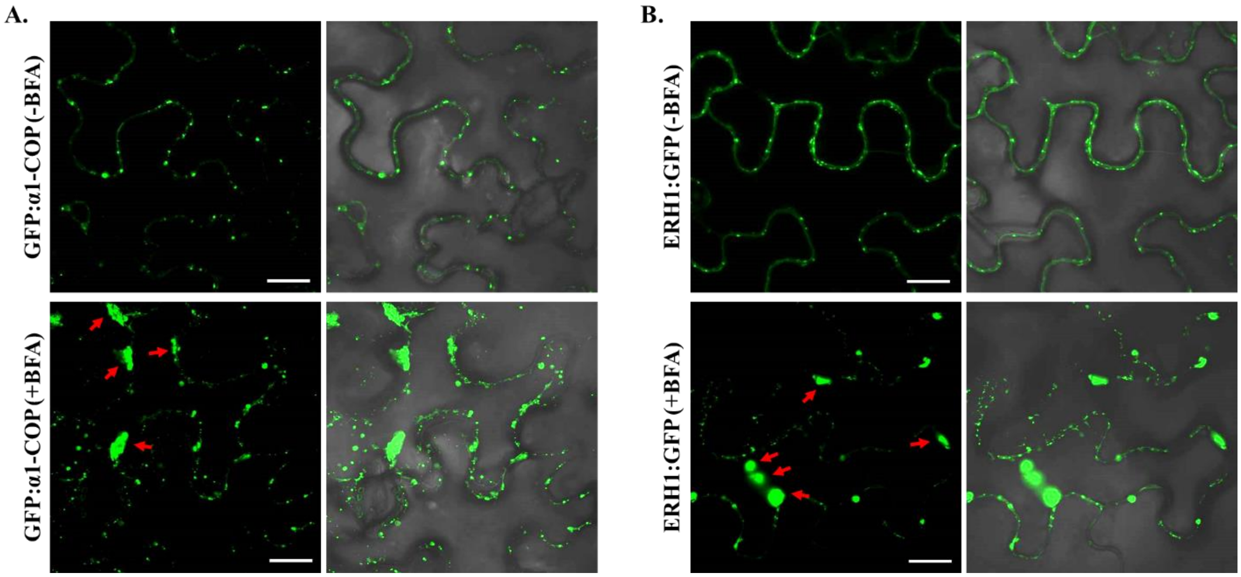


**Supplemental Figure 7. BFA disrupts intracellular transport of α1-COP and ERH1 proteins.**

**(A)** Confocal images of *N. benthamiana* transiently expressing GFP-α1-COP in the absence/presence of BFA (5 μg/ml of BFA was infiltrated into *N. benthamiana* leaves at 6 h prior to the observation). Red arrows indicate BFA compartments. Scale bars: 20 μm.

**(B)** Confocal images of *N. benthamiana* transiently expressing ERH1-GFP in the absence/presence of BFA (5 μg/ml of BFA was infiltrated into *N. benthamiana* leaves at 6 h prior to the observation). Red arrows indicate BFA compartments. Scale bars: 20 μm.

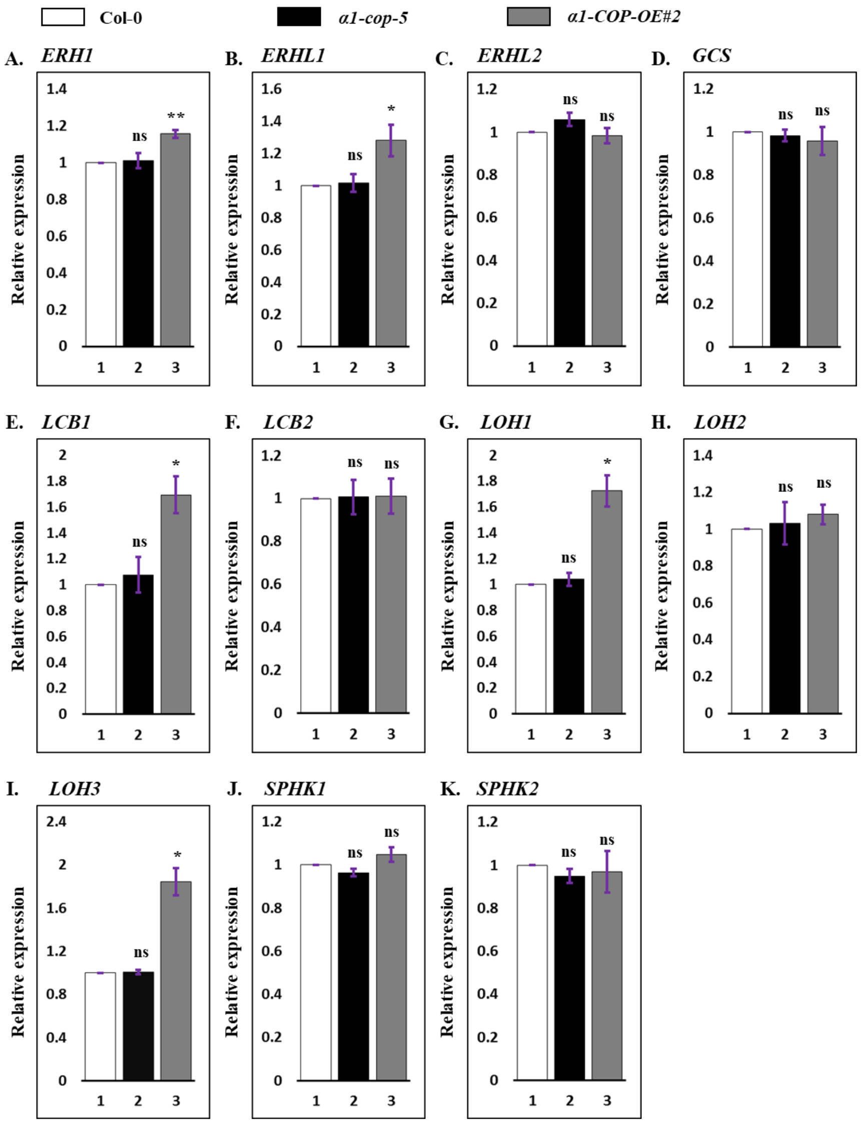


**Supplemental Figure 8. Quantitative RT-PCR analyses of sphingolipid enzymes in the wild-type Col-0, *α1-cop-5* mutant and α1-COP-OE#2 overexpression plants.**

Quantitative RT-PCR analyses of *ERH1* **(A),** *ERHL1* **(B)**, *ERHL2* **(C)**, *GCS* **(D)**, *LCB1* **(E)**, *LCB2* **(F)**, *LOH1* **(G)**, *LOH2* **(H)**, *LOH3* **(I)**, *SPHK1* **(J)**, *SPHK2* **(K)**, transcripts after normalization against four reference transcripts (*Actin, UBQ10, UBC9* and *EF-1α*) in total RNA samples of Arabidopsis wild-type Col-0 seedlings or *α1-cop-5* seedlings or α1-COP-OE2 seedlings. Bar graphs represent the average values of individual plant seedlings 10-day-old (*n*=25 for wild-type Col-0, *α1-cop-5* and α1-COP-OE#2). The Q-RT-PCR data represent mean values of two independent biological experiments, with four technical replicates. ±S.D. t-test (p-value of significant different classes are as follow: *, $p<0.05$; **, $p<0.01$ and ns means not significant).

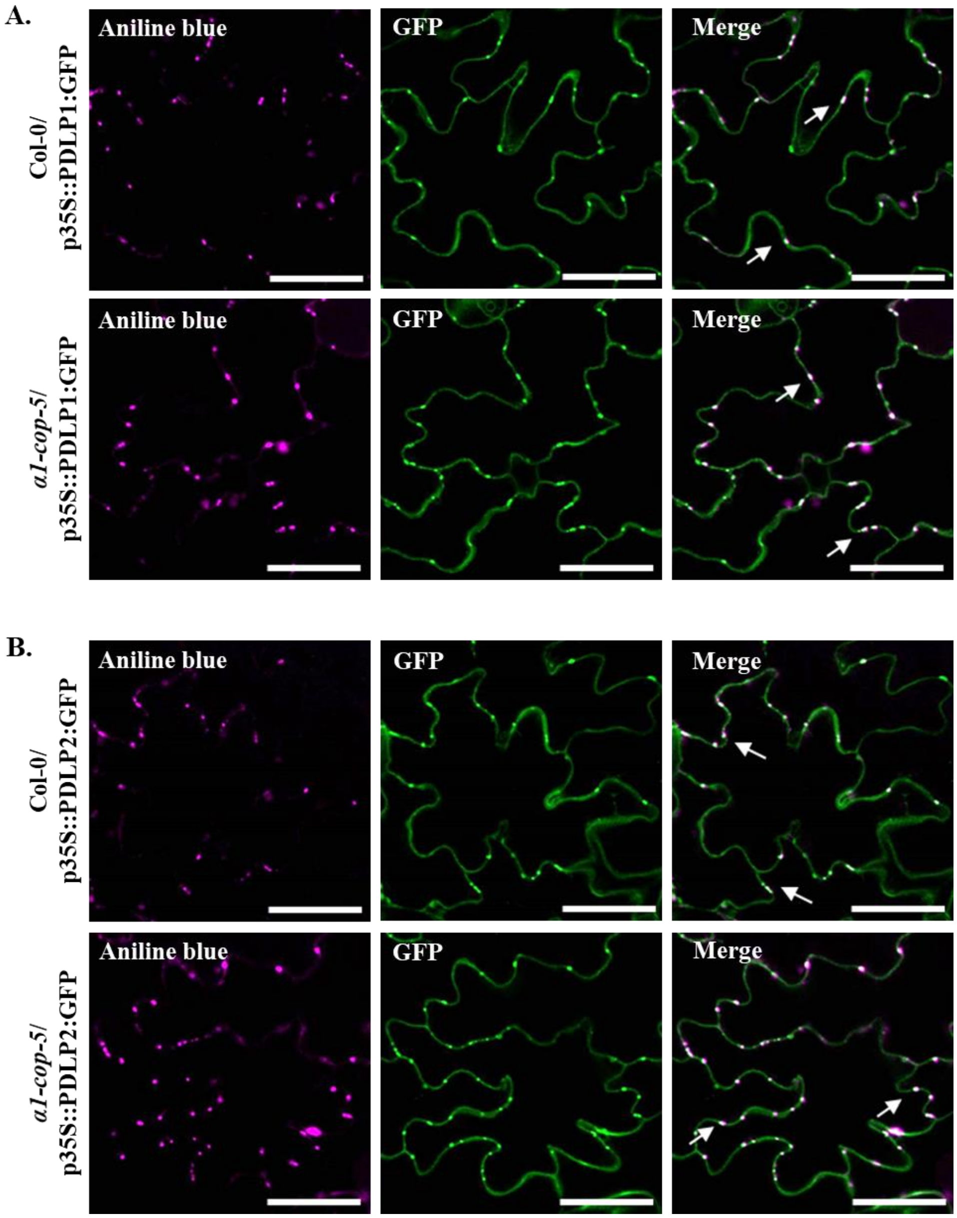


**Supplemental Figure 9. Subcellular localization of PDLP(s) proteins in the *α1-cop-5* mutant.** **(A)** Transgenic expression of PDLP1-GFP in Arabidopsis rosette leaves of wild-type Col-0 and *α1-cop-5* plants. The same samples were stained with aniline blue. White arrows indicate the GFP signals (green) co-localized with aniline blue signals (magenta) at PD. Scale bars: 50 µm. **(B)** Transgenic expression of PDLP2-GFP in Arabidopsis rosette leaves of wild-type Col-0 and *α1-cop-5* plants. The same sample were stained with aniline blue. White arrows indicate the GFP signals (green) co-localized with aniline blue signals (magenta) at PD. Scale bars: 50 µm.

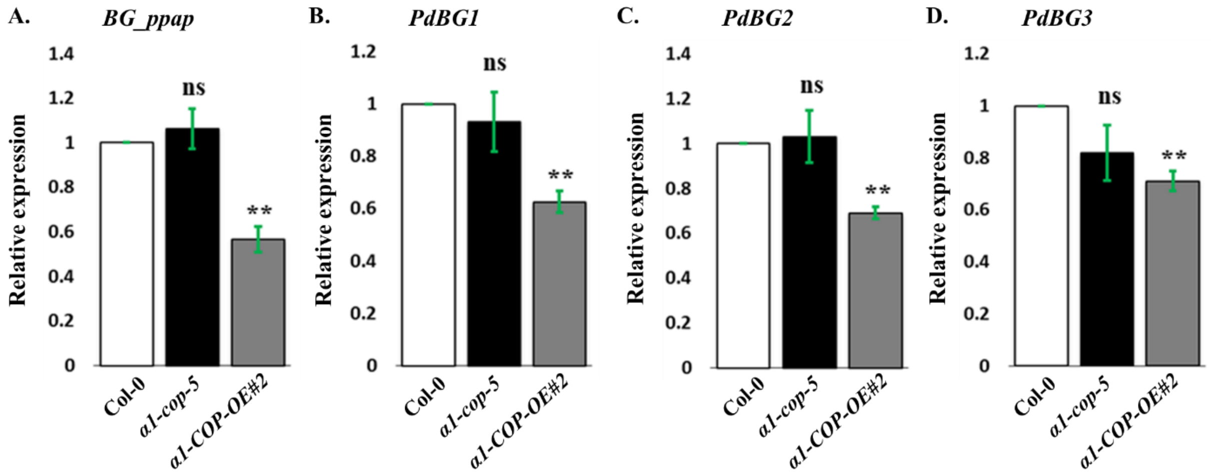


**Supplemental Figure 10. Quantitative RT-PCR analyses of callose degradation enzymes in wild-type Col-0, *α1-cop-5* mutant and α1-COP-OE#2 plants.**

Quantitative RT-PCR analyses of *BG_ppap* **(A)**, *PdBG1* **(B)**, *PdBG2* **(C)** and *PdBG3* **(D)** transcripts after normalization against four reference transcripts (*Actin, UBQ10, UBC9* and *EF-1α*) in total RNA samples of Arabidopsis wild-type Col-0, *α1-cop-5* and α1-COP-OE2 seedlings. Bar graphs represent the average values of individual plant seedlings 10-day-old (*n*=25 for wild-type Col-0, *α1-cop-5* and α1-COP-OE2). The Q-RT-PCR data represent mean values of two independent biological experiments, with four technical replicates. ±S.D. t-test (p-value of significant different classes are as follow: *, $p<0.05$; **, $p<0.01$ and ***, $p<0.001$, ns means not significant).

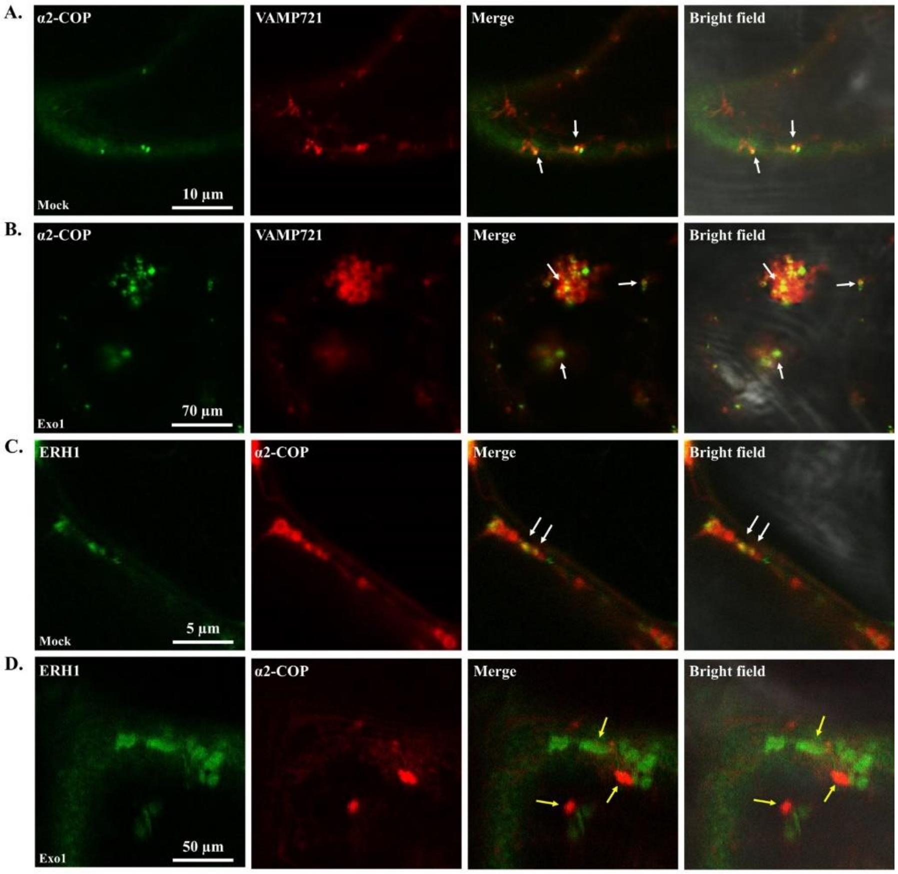


**Supplemental Figure 11. Subcellular localization of α2-COP and ERH1 in the presence of Exo1.**

**(A)** Confocal images of *N. benthamiana* epidermal cells transiently expressing GFP-α2-COP and VAMP721-RFP in mock condition. White arrows indicate co-localization of GFP and RFP signals. Scale bar: 10 µm.

**(B)** Confocal images of *N. benthamiana* epidermal cells transiently expressing GFP-α2-COP and VAMP721-RFP in the presence of Exo1. White arrows indicate co-localization of GFP and RFP signals. Scale bar: 70 µm.

**(C)** Confocal images of *N. benthamiana* epidermal cells transiently expressing ERH1-GFP and α2-COP-RFP in mock condition. White arrows indicate co-localization of GFP and RFP signals. Scale bar: 5 µm.

**(D)** Confocal images of *N. benthamiana* epidermal cells transiently expressing ERH1-GFP and α2-COP-RFP in the presence of Exo1. Yellow arrows indicate GFP and RFP signals do not co-localize. Scale bar: 50 µm.

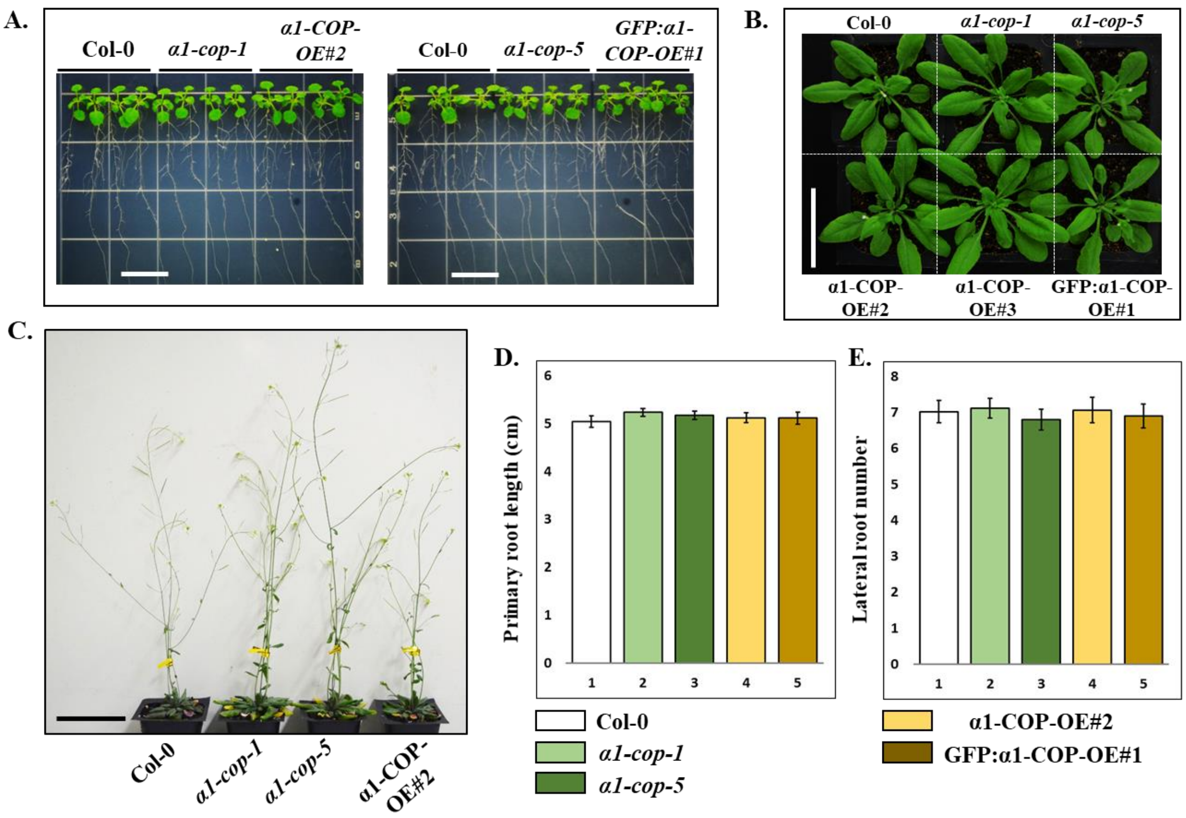


**Supplemental Figure 12. Growth of wild-type Col-0, *α1-cop* mutants and α1-COP overexpression plants.**

**(A)** Phenotypes of wild-type Col-0, *α1-cop* and α1-COP overexpression plants grown on normal MS medium. Scale bars: 1.5 cm.

**(B)** Phenotype of wild-type Col-0, *α1-cop* and α1-COP overexpression plants grown on the soil. Representative images were taken after 4 week grown on the soil. Scale bar: 5 cm.

**(C)** Phenotypes of wild-type Col-0, *α1-cop* and α1-COP overexpression plants grown on the soil. Representative images were taken after 6 week grown on the soil. Scale bar: 6 cm.

**(D)** Primary root length measurement of wild-type Col-0, *α1-cop* and α1-COP overexpression plants. Two-week-old seedlings were designated for primary root length measurement. Bar graphs represent the average values of individual plant seedlings (*n*=50).

**(E)** Lateral root number measurement of wild-type Col-0, *α1-cop* and α1-COP overexpression plants. Two-week-old seedlings were designated for lateral root number measurement. Bar graphs represent the average values of individual plant seedlings (*n*=50).

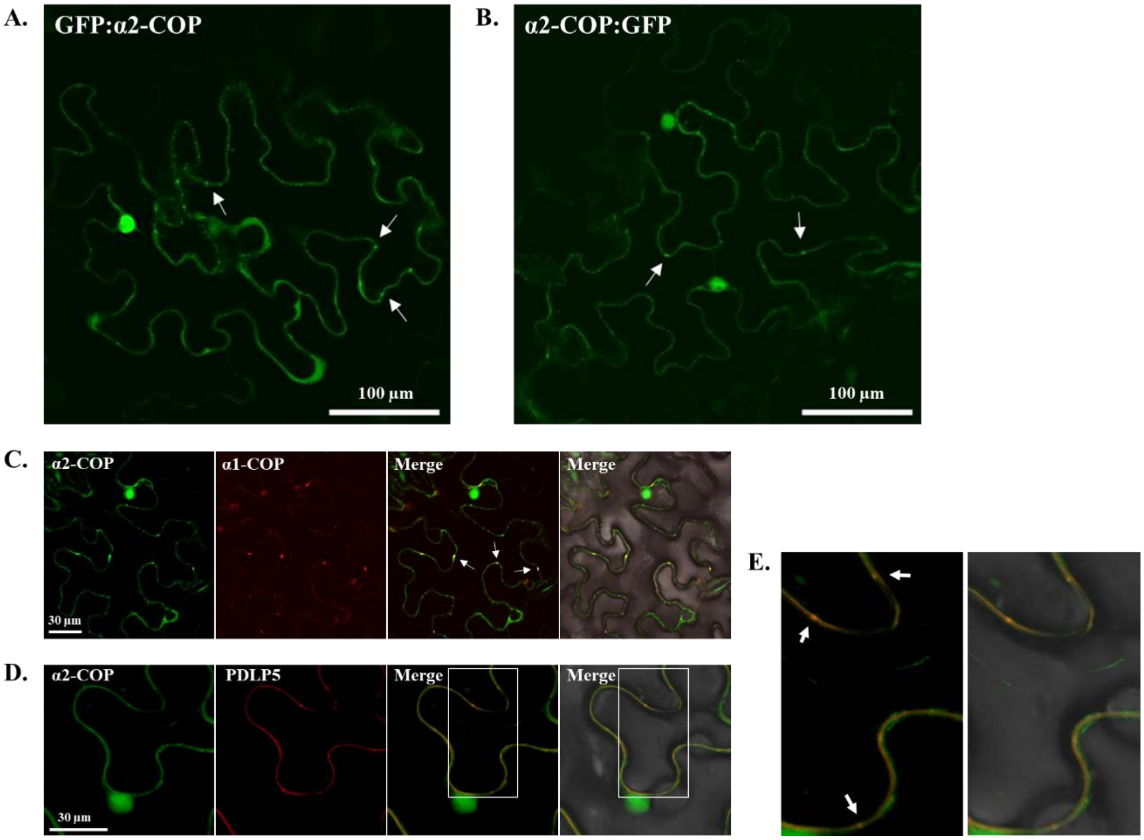


**Supplemental Figure 13. Subcellular localization of α2-COP protein.**

**(A)** Confocal image of *N. benthamiana* epidermal cells transiently expressing fusion protein of GFP-α2-COP. White arrows indicate punctate spots on the cell periphery. Scale bars: 100 μm.

**(B)** Confocal image of *N. benthamiana* epidermal cells transiently expressing fusion proteins of α2-COP-GFP. White arrows indicate punctate spots on the cell periphery. Scale bars: 100 μm.

**(C)** Confocal images of *N. benthamiana* epidermal cells transiently expressing GFP-α2-COP and α1-COP-RFP. α2-COP is partially colocalized with α1-COP (shown by white arrows). Scale bars: 30 μm.

**(D)** Confocal images of *N. benthamiana* epidermal cells transiently expressing GFP-α2-COP and PDLP5-RFP. α2-COP is partially colocalized with PDLP5 at PD. Scale bars: 30 μm.

**(E)** Co-localization of α2-COP and PDLP5 proteins depicted in **D** (white boxes). White arrows indicate that α2-COP and PDLP5 is co-localized at PD.

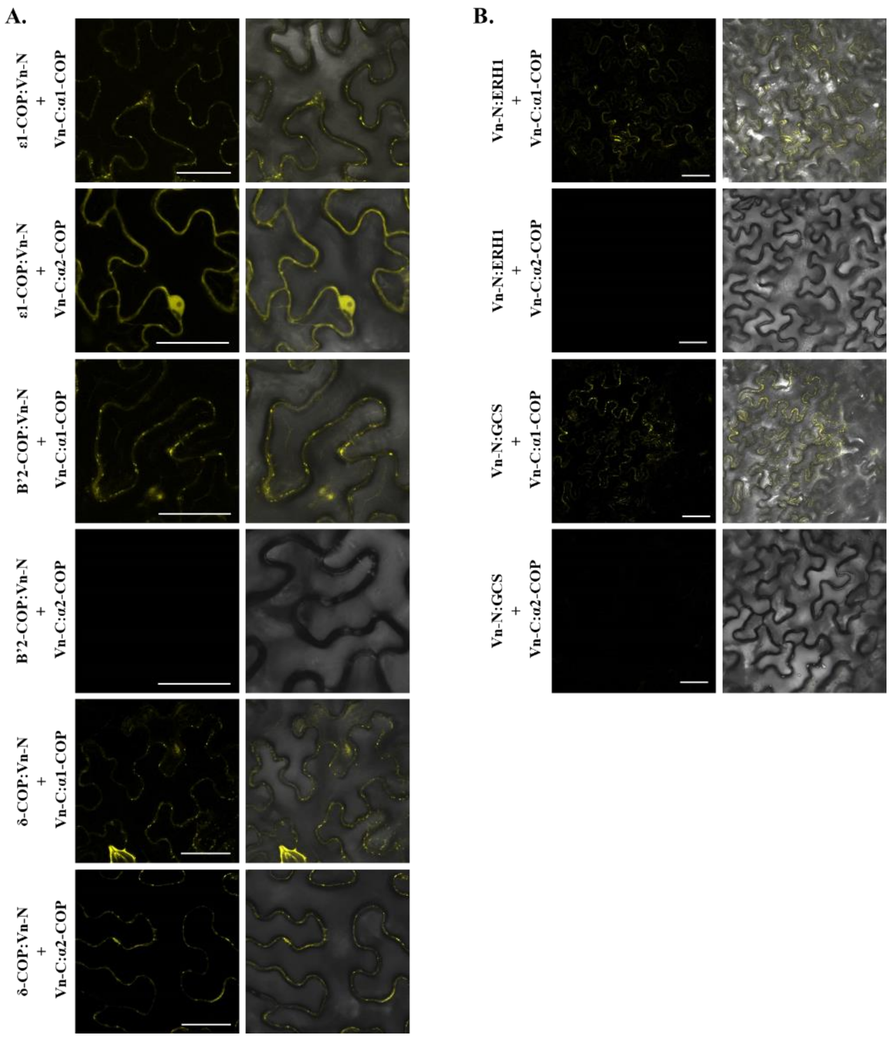


**Supplemental Figure 14. The interaction partner analysis of α2-COP protein with COPI sub-complex members and sphingolipid modifier enzymes.**

**(A)** BiFC analysis of α2-COP interaction with ε1-COP, β'2-COP and δ-COP in *planta*. Fluorescence was observed in the transformed *N. benthamiana* leaf epidermal cells, which results from complementation of the N-terminal part of the Venus fused with ε1-COP, β'2-COP and δ-COP by the C-terminal part of the Venus fused with α2-COP. The combinations between α1-COP, ε1-COP, β'2-COP and δ-COP were used as positive control. Scale bars: 50 µm.

**(B)** BiFC analysis of α2-COP interaction with ERH1 and GCS in *planta*. Fluorescence was observed in the transformed *N. benthamiana* leaf epidermal cells, which results from complementation of the N-terminal part of the Venus fused with ERH1 and GCS by the C-terminal part of the Venus fused with α2-COP. The combinations between α1-COP, ERH1 and GCS were used as positive control. Scale bars: 50 µm.
